# Single-cell landscape of nuclear configuration and gene expression during stem cell differentiation and X inactivation

**DOI:** 10.1101/2020.11.20.390765

**Authors:** Giancarlo Bonora, Vijay Ramani, Ritambhara Singh, He Fang, Dana Jackson, Sanjay Srivatsan, Ruolan Qiu, Choli Lee, Cole Trapnell, Jay Shendure, Zhijun Duan, Xinxian Deng, William S. Noble, Christine M. Disteche

## Abstract

Mammalian development is associated with extensive changes in gene expression, chromatin accessibility, and nuclear structure. Here, we follow such changes associated with mouse embryonic stem cell differentiation and X inactivation by integrating, for the first time, allele-specific data obtained by high-throughput single-cell RNA-seq, ATAC-seq, and Hi-C. In differentiated cells, contact decay profiles, which clearly distinguish the active and inactive X chromosomes, reveal loss of the inactive X-specific structure at mitosis followed by a rapid reappearance, suggesting a ‘bookkeeping’ mechanism. In differentiating embryonic stem cells, changes in contact decay profiles are detected in parallel on both the X chromosomes and autosomes, suggesting profound simultaneous reorganization. The onset of the inactive X-specific structure in single cells is notably delayed relative to that of gene silencing, consistent with the idea that chromatin compaction is a late event of X inactivation. Novel computational approaches to effectively align single-cell gene expression, chromatin accessibility, and 3D chromosome structure reveal that long-range structural changes to chromosomes appear as discrete events, unlike progressive changes in gene expression and chromatin accessibility.

## Introduction

Embryonic development is associated with genome-wide changes in gene expression, including silencing of pluripotent genes and onset of gene expression specific to various cell lineages. Global changes in gene expression are correlated with a drastic remodeling of the epigenetic landscape of chromatin, changes in DNA replication timing, and reorganization of the three-dimensional (3D) nuclear structure (Bonev et al. 2017; Du et al. 2017; Ke et al. 2017; Collombet et al. 2020; Miura et al. 2019). While studies have followed 3D structure remodeling events during development, most have been focused on very early developmental stages after fertilization and few have addressed events at the chromosomal scale in single cells.

Mammalian X chromosome inactivation (XCI), which randomly silences one of the two X chromosomes in early female embryogenesis, epitomizes a chromosome-wide change in gene expression and 3D structure, and takes place during early development (Lyon 1961). XCI is initiated by *Xist*, a long non-coding RNA (lncRNA) that coats *in cis* the future inactive X chromosome (Xi) and recruits/removes specific proteins or protein modifications to silence X-linked genes and condense the Xi into a heterochromatic structure (Galupa and Heard 2018; Giancarlo Bonora and Disteche 2017). Studies using sequencing-based assays and imaging to follow *in vitro* differentiation of female mouse embryonic stem cells (ESCs) provide a roadmap of the dynamics of gene silencing and structural organization of the Xi during the onset and establishment of XCI (Brockdorff, Bowness, and Wei 2020). *Xist* RNA interacts with RNA-binding proteins and master structural proteins, which help reshape one of the two X chromosomes into a unique condensed bipartite structure (Galupa and Heard 2018; Dossin et al. 2020; Brockdorff, Bowness, and Wei 2020). In both human and mouse, the two superdomains of long-range chromatin interactions are separated by a hinge region containing the conserved lncRNA locus *DXZ4/Dxz4* that represents a structural platform for frequent long-range contacts with multiple X-linked loci (Rao et al. 2014; Darrow et al. 2016; Giorgetti et al. 2016; G. Bonora et al. 2018; X. Deng et al. 2015; Minajigi et al. 2015; Wang et al. 2018). Once established, gene silencing and condensation of the Xi bipartite structure are stably maintained in somatic cells. However, studies of Xi-specific epigenetic and structural features point to mechanistic differences between establishment and maintenance of XCI (Giorgetti et al. 2016; Wang et al. 2018; Gdula et al. 2019; Wang et al. 2019; Jansz et al. 2018; Splinter et al. 2011; Marks et al. 2009; Fang et al., n.d.).

Our understanding of molecular and structural events associated with embryonic development, cell differentiation, and XCI is heavily based on ensemble datasets derived from bulk cells or tissues, which fail to address cell-to-cell dynamics of 3D structure and gene expression regulation, especially in relation to the cell cycle and stem cell differentiation (Bonev et al. 2017; Du et al. 2017; Ke et al. 2017; Keniry and Blewitt 2018; Miura et al. 2019; Froberg et al. 2018). Cell-to-cell variability in allelic gene expression has been reported based on single-cell RNA-seq analyses in mouse and human systems in which alleles can be distinguished (Borensztein et al. 2017; Q. Deng et al. 2014; Chen et al. 2016; Reinius et al. 2016; Cheng et al. 2019). Furthermore, super-resolution microscopy and single-cell Hi-C reveal variable 3D structures among nuclei, in part reflecting cell cycle changes (Nagano et al. 2013, 2017; Z. Liu et al. 2017; Bintu et al. 2018; Ramani et al. 2017; J. Liu et al. 2018; Finn et al. 2019; Cattoni et al. 2017; Flyamer et al. 2017; Stevens et al. 2017; Carstens, Nilges, and Habeck 2016; Tan et al. 2018; Su et al. 2020). Hence, methods have been developed to analyze single-cell Hi-C data and extract topics to accurately classify cell types in spite of variability among individual cells (Kim et al. 2020). Single-cell omics, especially Hi-C, have not been systematically applied to stages of ESC differentiation for a comprehensive integration of allele-specific data on gene expression, chromatin accessibility and 3D nuclear structure.

To gain insights into cellular heterogeneity and allelic dynamics in the structure and gene expression landscape of the genome during ESC differentiation and during XCI, we applied a single-cell combinatorial indexed (sci) approach for Hi-C, RNA-seq, and ATAC-seq (Ramani et al. 2017; Cao et al. 2017; Cusanovich et al. 2018). These sci-omics analyses were carried out in parallel in a time course of differentiation of female and male hybrid mouse ESCs into embryoid bodies (EBs) and in two differentiated hybrid cell lines with stable and completed XCI, embryonic kidney-derived Patski fibroblasts and EB-derived neural precursor cells (NPCs). Integration of datasets that measure large-scale chromosome structure together with gene expression and chromatin accessibility in thousands of single cells was performed to evaluate concurrent changes during ESC differentiation.

## Results

### 1. The 3D structure of the Xi detected by sci-Hi-C changes through the cell cycle

We initially focused our analysis on the sci-Hi-C data from Patski fibroblasts that contain a stable Xi from C57BL/6J (B6) and an active X chromosome (Xa) from *Mus spretus* (*spret*), which facilitates allelic studies (Fig. 1). Sci-Hi-C data were obtained from a total of 4,493 cells with a median coverage of 5,167 informative contacts (inter- and intrachromosomal contacts >1 kb) per cell, resulting in about 26M allelic contacts throughout the genome and 1.3M allelic contacts resolved on the Xi and the Xa (Supplementary Table 1). Previous bulk Hi-C studies in this cell line have demonstrated that the Xi exhibits a unique bipartite structure with two superdomains, whereas the Xa has a A/B compartment and TAD structures similar to that of autosomes (X. Deng et al. 2015; G. Bonora et al. 2018). Pseudobulk allelic contact maps for the X chromosome (chrX) and chromosome 1 (chr1) homologs using contacts aggregated from the 4,493 Patksi cells (Fig. 2A, E) or from subsets of cells (Extended Data Fig. 1A, B) are virtually identical to contact maps obtained from bulk data.

**Figure 1.**
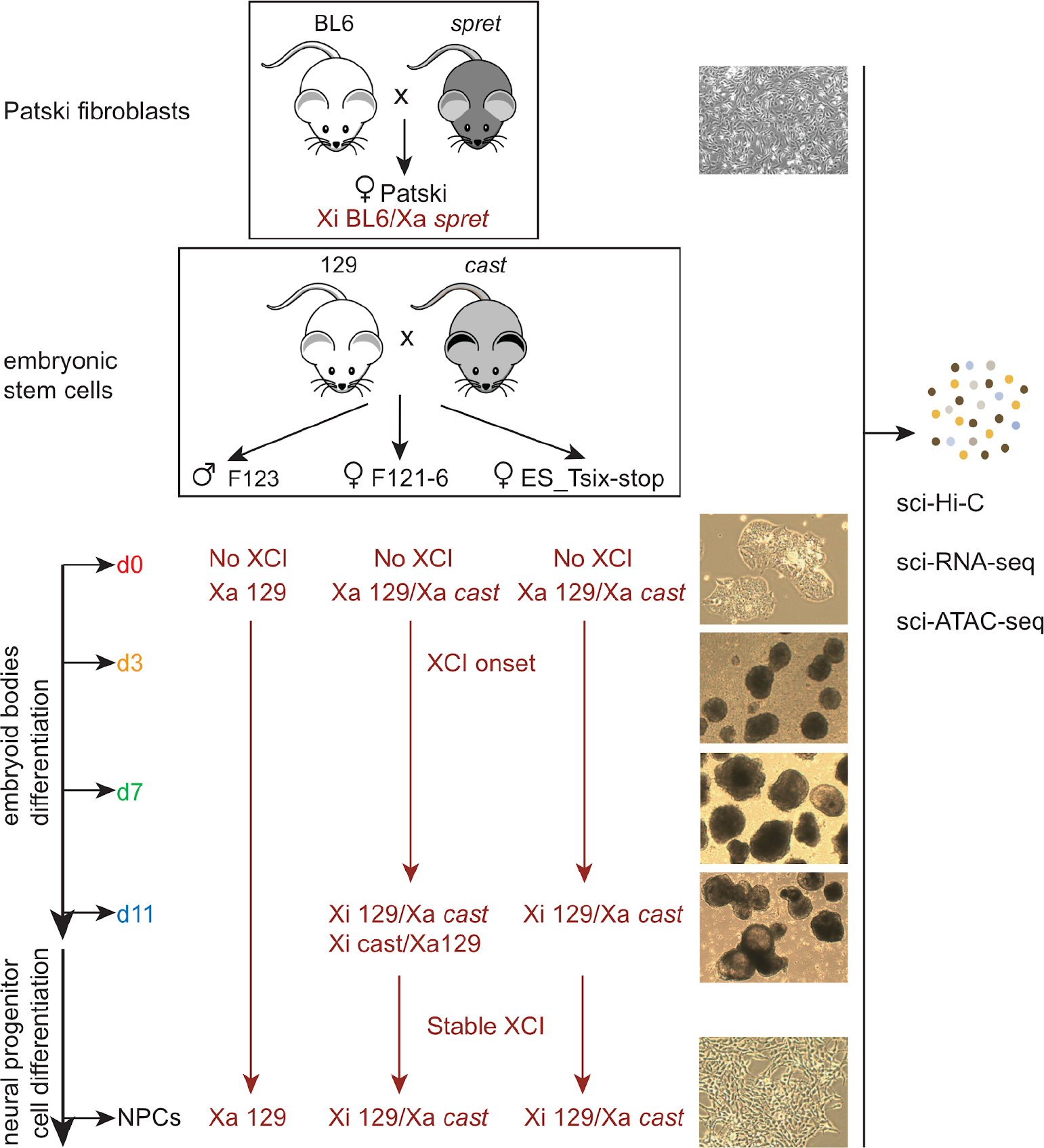
Schematic of the mouse cell lines used to generate sci-Hi-C, sci-RNA-seq, and sci-ATAC-seq. A B6 x *spret* cross was used to generate Patski fibroblasts (top box), and 129 x *cast* crosses were used to generate mESCs F123 (male), F121-6 (female), and ES_Tsix-stop (female) (lower box). Cells were collected during mESC differentiation to embryoid bodies at d0, d3, d7, and d11, followed by collection of neural precursor cells (NPCs). The XCI status is indicated in dark red. The B6 chrX is always the Xi in Patski cells. The male F123 line has a single Xa from 129. The *cast* or 129 chrX can be inactivated at random in differentiated F121-6, but due to a strong *Xce* allele the 129 chrX is more often the Xi. In the cloned F121-6 NPC line the 129 chrX is always the Xi. In ES_Tsix-stop cells in which XCI has occurred, the 129 chrX is always the Xi. Images of each culture are shown at right. Single cells were processed for sci-Hi-C, sci-RNA-seq, and sci-ATAC-seq as described in the text.

**Figure 2.**
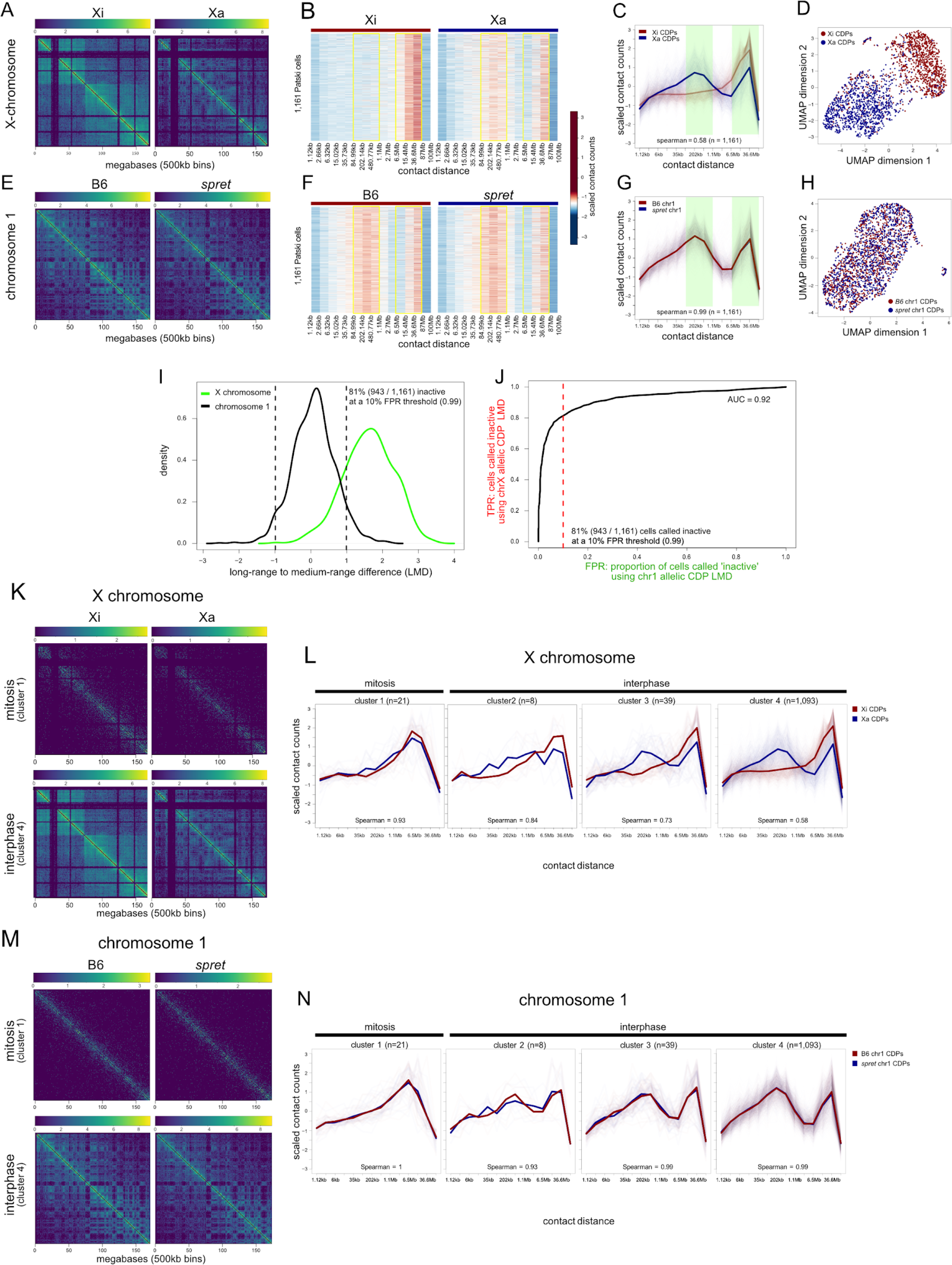
The Xi bipartite structure is captured by allelic sci-Hi-C data and changes with the cell cycle. **A)** Aggregate (pseudobulk) allelic contact maps for the Xi and Xa using all segregated contact read pairs from sci-Hi-C on 4,493 individual Patksi cells. Color scale in log10 (contact counts + 1). **B)** Heatmaps of allelic contact decay profiles (CDPs) for the Xi (red bar) and Xa (blue bar) for 1,161 Patski cells with at least 100 contacts per allele (within both chrX and chr1). The heatmap color scale reflects z-scaled counts within contact distance ranges shown along the x-axis. Contact counts were binned within exponentially-increasing contact distances ranges (2^x where x was incremented by 0.125). These bins were then aggregated further to reduce noise by combining the counts within 10 non-overlapping bins at a time. Yellow rectangles highlight the bins used for long-range to medium-range difference (LMD) calculation (see I, J and Methods). **C)** Plots of CDPs for the Xi (red) and Xi (blue) across the 1,161 cells in B. The average CDP is plotted over the plots of CDPs for each individual cell. The Spearman correlation between the average allelic CDPs is shown at the bottom of each plot. Contact counts were binned as described in B, and the bins used for long-range to medium-range difference (LMD) calculation are highlighted in green (see I, J and Methods). **D)** UMAP projections of CDPs for the Xi (red) and Xa (blue) for each of the 1,161 cells in B. Contact counts were binned as described in B. The CDPs for each parental allele form two distinct clusters. **E – H)** As in A – D, but for the B6 and *spret* (*Sp*) parental alleles of exemplar autosome, chr1. **I)** Distributions of the difference between the long-range to mid-range differences (LMD) between the allelic CDPs for each of 1,161 cells for chrX (green; as described in B) and chr1 (black; as described in F). The dashed line represents a threshold chosen below which cells were called as having an Xi bipartite structure based on the allelic CDP LMD. This threshold represents a 10% false positive rate (FPR) based on the distribution for chr1 allelic CDP LMD (see J). **J)** Plot of true positive rate (TPR; proportion of cells called called as having an Xi bipartite structure based on the LMD differences between allelic CDPs as described in B and I) versus false positive rate (FPR; proportion of cells called ‘inactive’ based on the LMD differences between the allelic CDPs for chr1 as described in F and I) at all thresholds of absolute LMD. The area under the curve (AUC) is 0.92. The dashed line represents the 10% FPR threshold chosen below which cells were called inactive based on the chrX allelic CDP LMD. **K)** Aggregate allelic contact maps for the Xi and Xa using contact pairs from 64 mitotic cells (top row) and 2,615 interphase cells (bottom row). Mitotic and interphase cells were grouped using the k-means clustering of the autosomal CDPs (Extended Data Fig. 3E). **L)** Plots of allelic CDPs for the Xi (red) and Xa (blue) for cells in each of the four clusters obtained from the k-means clustering of the autosomal CDPs (described in Extended Data Fig. 3E). The average CDP is plotted over plots of the CDPs for each individual cell. The spearman correlations between the average allelic CDPs is given at the bottom of the plot. **M – N)** As in K – L, but for maternal and paternal alleles of chr1.

The Xi shows a very distinctive contact decay profile (CDP) compared to the Xa or to the autosomes represented by the exemplar chr1 (Fig. 2B, F and Extended Data Fig. 2A, B). Heatmaps of allelic CDPs binned within exponentially-increasing contact distances ranges were plotted for each of 1,161 Patski cells with sufficient allelic coverage (Extended Data Fig. 2A, B and see Methods). As expected, a unique enrichment in long-range (6.5 Mb – 87 Mb) contacts was observed on the Xi compared to the Xa or to the two homologs of chr1 (Fig. 2B, F). This trend was accompanied by a reduction in mid-range contacts (85 kb – 1.1 Mb), consistent with much weaker TADs on the Xi (G. Bonora et al. 2018). These patterns were remarkably consistent across individual cells (Fig. 2B, F), suggesting a stable maintenance of the Xi structure. Plots of average CDPs versus contact distance clearly differentiate the Xi from the Xa, the latter resembling chr1 homologs, consistent with classic contact profiles reported for autosomes (Nagano et al. 2017) (Fig. 2C, G). UMAP projections of CDPs based on sci-Hi-C data clearly show a separation between the Xi from the Xa, except in a few cells in which there is an apparent lack of the characteristic Xi-specific structure, whereas, in contrast, projections CDPs for chr1 homologs show no separation (Fig. 2D, H).

Next, we defined parameters to call single cells as having an X chromosome with a bipartite condensed structure (hereafter, “bipartite X”), by plotting the distributions of Spearman correlations between the CDPs of each allele for chrX and chr1 for each of the selected 1,161 cells. This resulted in a clear separation between chrX and chr1: the chrX-allelic correlation in each cell is variable and low, resulting in a wide distribution that peaks close to 0.5, while the chr1-allelic correlation is consistent among cells, resulting in a sharp distribution that peaks close to 1 (Extended Data Fig. 2C). We then defined a threshold representing a 10% false positive rate (FPR) of calling a cell containing a bipartite X based on the distribution for chr1-allelic CDPs (Extended Data Fig. 2C, D). Using these parameters, a majority of Patski cells (79% or 917/1,161) were deemed to contain a bipartite X, supporting that CDP correlation analysis can be reliably used to ascertain the 3D structure status of the two X chromosomes in single cells. However, the correlation does not provide information about which allele has changed. We can assume that it is the B6 allele with the bipartite X, since XCI is skewed towards that allele in Patski cells. To confirm this, we assessed the difference between the total long-range (6.5 Mb – 87 Mb) and the total mid-range (85 kb – 1.1 Mb) contact counts, or the the long-range to mid-range difference (LMD; see Methods), for each homolog of each cell. If two chrX homologs display a significantly large difference between their LMDs, then the homolog with the larger LMD likely possesses the Xi 3D structure. Again we defined a threshold representing a 10% FPR for calling a cell containing a bipartite chromosome based on the distribution of LMD differences for chr1-allelic CDPs (Fig. 2I, J). Using these parameters, a majority of Patski cells (81% or 943/1,161 cells) were deemed to contain a B6 bipartite X, which is a similar proportion to that from the CDP correlation analysis. Indeed, 89% of Patski cells called inactive using the Spearman correlation between the CDPs of the homologs were also called inactive using the LMD approach. Thus, the results of the LMD analysis correspond closely to those obtained using CDP correlation analysis, with the additional benefit that they can be reliably used to ascertain which chrX homolog has assumed the Xi 3D structure.

The proportion of Patski cells lacking a Xi-bipartite call, as ascertained by LMD analysis, is dependent on setting the FPR threshold, but does not appear to represent cells that have lost the Xi since at least 100 B6-specific X-linked contacts were detected in these cells (see Methods). Furthermore, we have previously shown that more than 96% of Patski cells retain one Xa and one Xi, using both DNA FISH with an X-paint probe and RNA FISH with an *Xist* probe (Bonora et al., 2018). Thus, cells that lack a CDP difference characteristic of the Xi (Fig. 2D, I, J and Extended Data Fig. 2C, D) likely represent a specific stage of the cell cycle when chromatin is reorganized (Naumova et al., 2013; Nagano et al., 2017), or may represent cells where data is too sparse to make a confident call as to the state of the chrX homologs based on allelic CDPs.

To address cell cycle changes in Xi structure, cells were ordered through the cell cycle using biallelic CDP distributions on autosomes (Nagano et al. 2017). Cells were grouped into four clusters using *k*-means clustering using the Spearman correlation distance between the cells’ CDPs (Extended Data Fig. 2E, F). Aggregated allelic contact maps for the Xi and Xa and for chr1 were markedly different when comparing 64 mitotic cells in cluster 1 (1.4% of 4,493 cells) to 2,615 interphase cells in cluster 4, with the mitotic cells displaying the well documented strong diagonal component and no compartmentalization due to condensin-mediated polymer self-looping (Fig. 2K, M) (Naumova et al. 2013; Abramo et al. 2019; H. Zhang et al. 2019; Nagano et al. 2017). We then examined cell-cycle phased CDP profiles for the selected 1,161 Patski cells. Plots of allelic CDPs for the Xi and Xa for each cell in each of the four clusters were markedly similar in mitotic cells but diverged at other stages (Fig. 2L). At mitosis, the profiles for chrX and chr1 were similar for all alleles, consistent with the uniform condensed structure of mitotic chromosomes (Fig. 2L, N). As visualized in a video assembled by pooling contact maps for the Xi and Xa and for chr1 grouped based on their cell cycle stage, the bipartite X disappears during mitosis and rapidly re-emerges during interphase, as does the A/B compartment structure of the Xa and autosomes (Extended Data Video 1).

We conclude that in differentiated Patski fibroblasts the bipartite X is easily detected in a majority of nuclei by analysis of the CDP profiles. A rapid structural change takes place at mitosis, due to tight condensation of chromosomes by the condensin machinery. Once removed at the return of interphase the bipartite organization of the Xi quickly resumes, suggesting a “bookmark memory” of that structure, potentially represented by the clustered CTCF binding at *Dxz4*, which specifically separates the Xi superdomains.

### 2. Cells with an Xi bipartite structure progressively increase during ESC differentiation

As shown in the previous section LMD analysis provides a reliable method to detect the Xi bipartite structure in Patski cells, which represent a differentiated state. To follow the dynamics of the establishment of the Xi bipartite structure during differentiation, sci-Hi-C was done on embryonic stem cells derived from 129 x *cast* F1 embryos (Fig. 1). As in Patski cells, alleles in F1 hybrid ESCs can be distinguished based on single nucleotide polymorphisms (SNPs). We examined two female ESC lines, ES_Tsix-stop and F121, collected at d0, d3, d7, and d11 of EB differentiation, and a control male ESC line, F123, collected at d0, as well as NPCs differentiated *in vitro* from all three cell lines (Fig. 1). In addition, sci-Hi-C was done on cells derived from an independent EB differentiation time course of F123, F121, and ES_Tsix-stop cells, collected at d0 and d7, and on d20 fibroblasts enriched from EB outgrowth (time course experiment 2; see Methods). These time points were selected based on previous bulk studies of mESC EB differentiation at stages corresponding to pre-XCI (d0), early-XCI (d3), mid-XCI (d7), late-XCI (d11), and post-XCI (d20 outgrowth fibroblasts and NPCs) (Simon et al. 2013; Pinter et al. 2012; Lin et al. 2007; Marks et al. 2009, 2015; Gendrel et al. 2012). The number of cells, contacts per cell, and contacts per allele (129 and *cast*) are listed in Supplementary Table 1.

We first examined the establishment of the bipartite X during EB (d0-11) and NPC differentiation of ES_Tsix-stop, in which the 129 strain-derived X is always the one to be silenced during differentiation and thus is easy to track (Fig. 3A, B and Extended Data Fig. 3A, B). Plots of the distribution of mean allelic CDPs show the progressive appearance of the bipartite structure of the 129 X during differentiation. There is evidence of an increase in long-range contacts in the emerging Xi and a decrease in mid-range contacts, resulting in an increase in LMD, which can be used to identify which chrX homolog acquires a bipartite structure, as was done in Patski cells (Fig. 2C, I and 3A, B). In contrast, allelic CDPs appear similar for autosomal homologs as exemplified by chr1 at each time point (Extended Data Fig. 3A). The progressive appearance of a characteristic Xi structure as shown by the mean CDP analysis of differentiating ESCs could be due to two factors: an increase in the proportion of cells with an Xi structure and a progressively stronger Xi structure in individual cells at each time point. To examine these factors, we applied the LMD analysis to classify cells with a bipartite X at each time point. At d0 the proportion of cells identified as having a bipartite 129 chrX was 9% (Fig. 3B, E and Supplementary Table 2). This proportion increased to 34%, 63%, 71%, and 83% at d3, d7, d11, and NPCs, respectively, consistent with each stage representing a mixture of cells with or without the Xi structure. Similar proportions of cells assuming a bipartite X were obtained using the Spearman correlation coefficient between allelic CDPs (Extended Data Fig. 3B).

**Figure 3.**
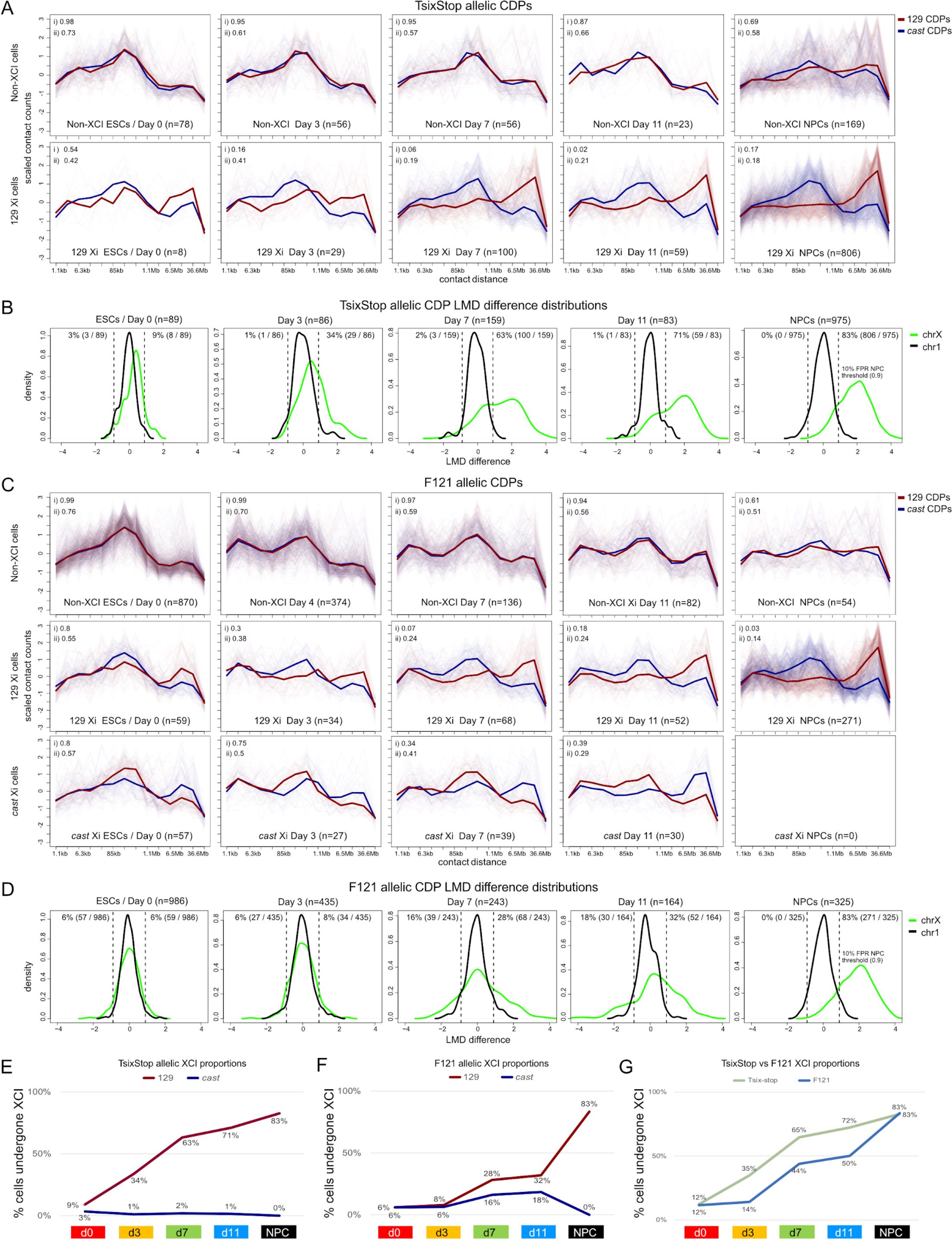
X inactivation-associated structural changes in cells during differentiation. **A)** CDP plots for chrX parental alleles (129 in red, *cast* in blue) in ES_Tsix-stop at the indicated time points during differentiation from ESCs (d0) to embryoid bodies (d11), as well as neural precursor cells (NPCs) for cells classified as not having the Xi bipartite structure (top row) and those that do exhibit CDPs indicative of an Xi on their 129 allele (bottom row; XCI is skewed towards the 129 allele in ES_Tsix-stop cells). Only cells with at least 50 contacts per allele within both chrX and chr1. The plots show z-scaled counts within contact distance ranges shown along the x-axis. Contact counts were binned within exponentially-increasing contact distances ranges (2^x where x was incremented by 0.125). These bins were then aggregated further to reduce noise by combining the counts within 10 non-overlapping bins at a time. Number of cells at each time point is given by the n number. The average allelic CDP is plotted over the plots of CDPs for each individual cell. Spearman correlation values between the two mean allelic CDPs (i) and median of the Spearman correlation values between allelic CDPs for each cell (ii) are given in the top left corner of each plot. **B)** Distributions of the difference between the long-range to mid-range differences (LMDs) of each allele (as described in Figure 2I and Methods) for each cell as described in A for chrX (green) and chr1 (black). The dashed lines represent a threshold chosen below which cells were called as having an Xi bipartite structure based on the chrX allelic CDP LMD differences. This threshold represents a 10% false positive rate (FPR) based on the distribution for chr1 allelic CDP LMD differences. See Supplementary Table 2. **C)** As in A, but for F121 cells with additional panels showing the allelic CDPs for those cells that do exhibit CDPs indicative of an Xi bipartite structure on their *cast* allele (bottom row; XCI is partially skewed towards the 129 allele in F121 cells at d0-11 and completely skewed to the 129 allele in NPCs). **D)** As in B, but for F121 cells. **E)** Plots of the proportion (%) of ES_Tsix-stop cells with a 129 allele (red) and *cast* (blue) allele with an Xi bipartite structure. **F)** As in E but for F121 cells. **G)** Plots of the proportion (%) of ES_Tsix-stop cells (light green) and F121 cells (light blue) with an Xi bipartite structure.

Next, we analyzed F121 cells, which undergo random XCI albeit with ∼3:1 skewing towards the 129 X due to a stronger X-controlling element (Xce) on the *cas*t allele (Cattanach and Rasberry 1994). Based on LMD analysis, the percentage of cells with a bipartite X was 12% (6% 129 X) at d0, and increased to 14% (8% 129 X), 44% (28% 129 X), and 50% (32% 129 X) at d3, d7, and d11, respectively, consistent with progressive selection of cells with a 129 Xi (Fig. 3C, D, F and Supplementary Table 2). Similar proportions of cells were obtained using Spearman correlation coefficients between allelic CDPs (Extended Data Fig. 3C, D). Note that at d0 the CDPs for both chrX and chr1 are quite different from those at later time points, with fewer short-range and more mid-range contacts (Extended Data Fig. 3C). As in Tsix-stop NPCs, XCI in cloned F121 NPCs is completely skewed towards the 129 allele, since this is a single-cell clonal line (Fig. 3D, F). Reassuringly, the proportion of cells assessed to have an Xi-specific structure is the same for the Tsix-stop and F121 NPCs (83%), and is also similar to that for Patski cells (81%). However, the proportion of F121 cells that acquire the Xi-specific structure increases more slowly than that of ES_Tsix-stop cells (Fig. 3G). This is probably because the 129 chrX carrying a *Tsix* mutation cannot repress upregulation of *Xist* upon differentiation, and thus is forced to undergo inactivation. A similarly slow XCI onset in normal female ESCs compared to the ES_Tsix-stop line was previously reported based on the proportion of cells acquiring an *Xist* cloud during differentiation, with the delay being ascribed to the process of stochastic choice (Monkhorst et al. 2008).

We further confirmed the appearance of a bipartite X in female cells by assembling a pseudobulk allelic contact heatmap for the presumptive Xi by aggregating contacts in a total of 433 F121 and ES_Tsix-stop cells deemed to have a bipartite X call at d3, d7 and d11. This clearly shows the emergence of the bipartite X, which is not seen for the corresponding Xa heatmap (Extended Data Fig. 4A). The bipartite X becomes more distinctive in the heatmap aggregated from the Xi contacts for 1,077 F121 and Tsix-stop NPCs that have a bipartite X call, and is associated with a pronounced dip in the contact score across the *Dxz4* locus (Extended Data Fig. 4B). Analysis of a sci-Hi-C dataset for an independent EB differentiation time course with comparatively fewer differentiating cells further confirmed the progressive appearance of cells with a bipartite X during differentiation and the efficacy of the LMD analysis (Extended Data Fig. 4C-F and Supplementary Table 2). We found that 59% of d20 ES_Tsix-stop fibroblasts (56% 129 Xi; 3% *cast* Xi) and 74% of F121 d20 fibroblasts (49% 129 Xi; 25% *cast* Xi) had a bipartite X call, consistent with their completely or partially skewed XCI patterns respectively. In addition, we examined ES (d0) cells and NPCs from the male F123 cell line. Allelic CDPs for chr1 homologs and for the single active X chromosome appear similar to those in F121 (Extended Data Fig. 3E).

In conclusion, the proportion of nuclei exhibiting a high frequency of ultra-long range interactions characteristic of the Xi-specific 3D structure increases progressively during the course of differentiation of female cells. While we cannot exclude the existence of intermediate structures undetectable by our approach our single cell data show that at each differentiation time point there is evidence of a mixture of cells with the bipartite X and cells without it, something that would not be evident from bulk sci-Hi-C data. Whether cells in different embryonic lineages acquire an Xi structure at different rates remains to be determined..

### 3. Allelic trajectory analysis reveals discrete changes to 3D genome organization

We next subjected genome-scale allelic CDP data (i.e. allelic CDPs for each chromosome concatenated together) to semi-supervised trajectory analysis using the implementation of the DDRtree algorithm (Mao et al. 2015) in Monocle2 (Qiu et al. 2017) based on differential CDP bins across time points. This yielded a bifurcated developmental trajectory dependent on the time points of differentiation, even though some cells from different time points were dispersed across all three branches, indicative of variability in nuclear structure (Fig. 4A). Notwithstanding this variability, most d0 cells (80%) fall along one of the three branches (called early branch), representing the root of the developmental trajectory, while most d3 cells (86%) and about half of d7 cells (45%) are found along a second (intermediate) branch seemingly representing a transitional stage of differentiation. The third (differentiated) branch comprises a majority of NPCs (77%), d11 cells (74%), and about half of d7 cells (46%), and thus appears to capture cells exhibiting a nuclear conformation representative of a more differentiated stage of development (Fig. 4A). Cells at the terminal end of the differentiated branch are later in pseudotime than those at the terminal end of the intermediate branch, which provides further evidence that the intermediate branch represents a conformation that arises during the course of differentiation (Extended Data Fig. 5A). A majority of cells (56-65%) in the trajectories cluster at the terminal ends of each branch, which suggests the existence of three distinct types of nuclear structure as defined by CDPs (Fig. 4A and Extended Data Fig. 5B). A similar trajectory with three branches was obtained when NPCs were omitted, which shows that the three conformational states captured by each branch represent structural changes that occur during differentiation and are not driven by a specifically differentiated cell type (Extended Data Fig. 5C). Similar trajectories were also obtained based on autosomal and X-chromosomal allelic CDPs, supporting profound 3D structural changes throughout the genome during differentiation (Fig. 4B, C and Extended Data Fig. 5E).

**Figure 4.**
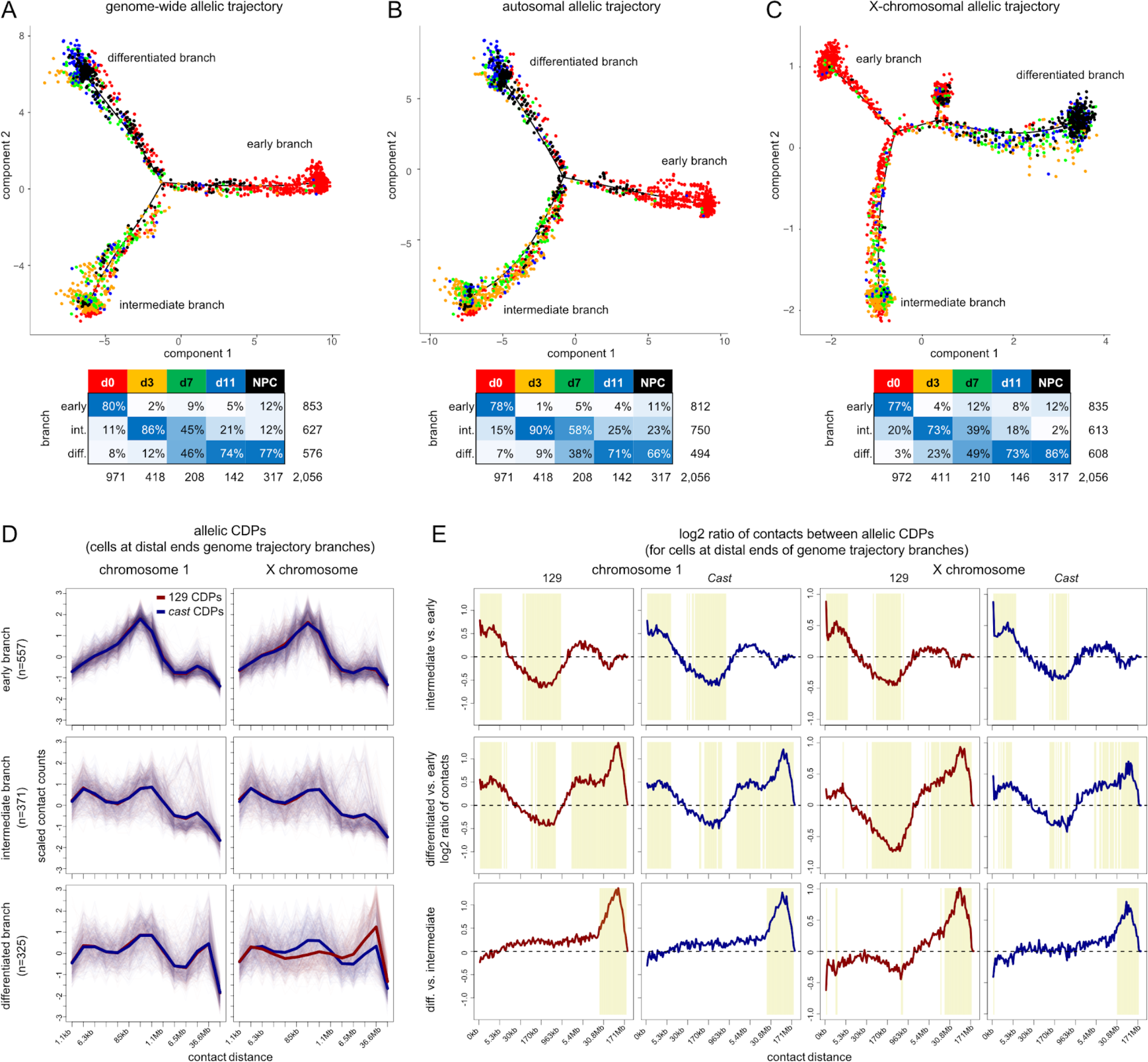
Trajectory analyses based on allelic CDPs reveal distinct conformational states during differentiation. **A)** Semi-supervised genome-wide trajectory analysis of F121 cells based on allelic CDPs (see Fig. 2I and Methods) from sci-Hi-C data using Monocle2 DDRTree. Allelic CDPs for each chromosome were concatenated together for each of 2,056 F121 cells with at least 100 contacts per chr1 and chrX allele. 2,040 significantly differential bins from the genome-scale concatenated allelic CDPs across time points with q-value <= 10^−11^ were used to construct the trajectory. Each data point represents a cell colored by time point. Arrows indicate the direction of development in pseudotime. Branches are labeled according to developmental state: branch 1 is the root embryonic stem cell (early) branch; branch 2 captures an intermediate state (intermediate branch); and branch 3 captures the genome architecture of differentiated cells including NPCs (differentiated branch). Inset table gives cell counts per time point per trajectory branch and time point color for data points in the plot. **B)** As in A, but based on 1,924 significantly differential bins (q-value <= 10^−11^) for autosomal allelic CDPs. **C)** As in A, but based on 178 significantly differential bins (q-value <= 10^−5^) for chrX CDPs. Three early sub-branches collapsed into a single early branch. **D)** Allelic CDPs for chr1 (left column) and chrX (right column) (129 in red, *cast* in blue) for cells at the distal ends of the three branches (rows) from the trajectory described in A (see Extended Data Fig. 6B). Binning and scaling details of the plots as described in Fig. 3A. **E)** Plots of the log2 ratios between allelic CDPs for cells at the distal ends of each pair of branches (rows) in the trajectory described in A (Extended Data Fig. 6B) for every logarithmic bin (2^x where x was incremented by 0.125) for the 129 (red) and *cast* (blue) alleles of chr1 and chrX. Top row: intermediate vs. early branch. Middle row: differentiated vs early branch. Bottom row: differentiated vs. intermediate branch. The dashed line reflects no difference between the two groups of cells and yellow highlighting reflects bins that show a significant difference (adjusted p-value <10^-6) between the two groups of cells (Extended Data Fig. 6D).

To identify structural features specifically associated with each of the three relatively discrete 3D conformations, the average CDPs for chr1 and chrX were derived from the cells at the end of each branch, regardless of their day of differentiation (Fig. 4D and Extended Data Fig. 5B). Comparisons between these average CDPs show a distinct distribution of contacts for both chr1 and chrX in cells within each of the three branches (Fig. 4E and Extended Data Fig. 5D). Cells in the early branch have significantly fewer very-short-range contacts (<10 kb) corresponding to chromatin loops and more medium-range contacts (30 kb – 1 Mb) corresponding to TAD/sub-TAD structures compared to those in either the intermediate or the differentiated branches. Cells in the intermediate and differentiated branches acquire very-long range contacts in the 30 – 170 Mb range, corresponding to the formation of A/B compartments. A comparison between the CDPs for cells in the intermediate and differentiated branches shows little change in short-range and medium-range contacts but a strong increase in very long-range contacts associated with differentiation. While chr1 homologs show no differences between alleles, there are strong differences between chrX alleles, with the *cast* Xa resembling chr1, and the 129 Xi showing unique structural changes due to the emergence of its bipartite structure (Fig. 4D, E). Both the decrease in medium-range contacts and the increase in very-long range contacts are exaggerated on the Xi in differentiated cells versus earlier stages, which likely reflects the loss of A/B compartments and the overall condensation of the Xi.

In conclusion, trajectory analyses of sci-Hi-C CDP data reveal three distinct nuclear structure states reflecting discrete and profound changes not only in the structure of the X chromosomes but also to that of autosomes. The order of events that take place during differentiation appears to be that short range contacts are gained and medium-range contacts lost before very long range contacts fully develop. The Xi appears to acquire its bipartite structure at the same time that autosomes show a sharp increase in very long-range contacts. Thus, our single cell Hi-C analysis shows for the first time that nuclei abruptly change structure during differentiation, which contrasts with the smoother linear developmental trajectories typically observed based on expression or accessibility data, as shown in the next section.

### 4. Single-cell expression and accessibility trajectories recapitulate ESC differentiation and XCI

Changes in gene expression and chromatin accessibility during differentiation of female F121 and male F123 cells were measured by sci-RNA-seq and sci-ATAC-seq applied to cells collected on d0, d3, d7, and d11, and after differentiation into NPCs (Fig. 1 and Supplementary Tables 3 and 4). With sci-RNA-seq, we obtained 9,021 cells with ≥200 UMIs (Extended Data Fig. 6A-C). By sci-RNA-seq median autosomal and X-linked gene expression was highest at d0 and in NPCs (Extended Data Fig. 6C). With sci-ATAC-seq we obtained 2,727 cells with ≥500 UMIs (Extended Data Fig. 6D-F).

A non-allelic developmental trajectory was obtained using the same DDRtree algorithm used in the previous section, but in this case based on differentially expressed genes in sci-RNA-seq data across the time points in both F121 and F123 cells (Fig. 5A). This semi-supervised trajectory has a single branch point representing the divergence of NPCs from the default EB differentiation pathway. The ESC branch mainly comprises d0 (79%) and some d3 cells (19%), while the EB branch mainly comprises d7 (43%) and d11 cells (38%), and no NPCs. A separate NPC-specific branch includes a majority of NPCs (82%) and some cells from earlier stages, especially d11 cells (9%). Developmental pseudotime can be assigned to each cell in the trajectory based on that cell’s distance from the root node – in this case, ESCs. Individual genes implicated in developmental processes demonstrate changes in expression as a function of pseudotime (Fig. 5B). For example, genes associated with pluripotency of ESCs (e.g. *Klf4*) show a decrease in expression during differentiation. *Otx2*, a homeobox gene important for early embryonic patterning of the brain and heart, is transiently expressed, while *Sema5a*, a gene encoding a bifunctional axon guidance cue for mammalian midbrain neurons, is gradually upregulated during differentiation, and *Map2*, an NPC marker gene, is specifically expressed in NPCs. As expected, *Xist* expression progressively increases only in female cells during differentiation, consistent with the onset and establishment of XCI.

**Figure 5.**
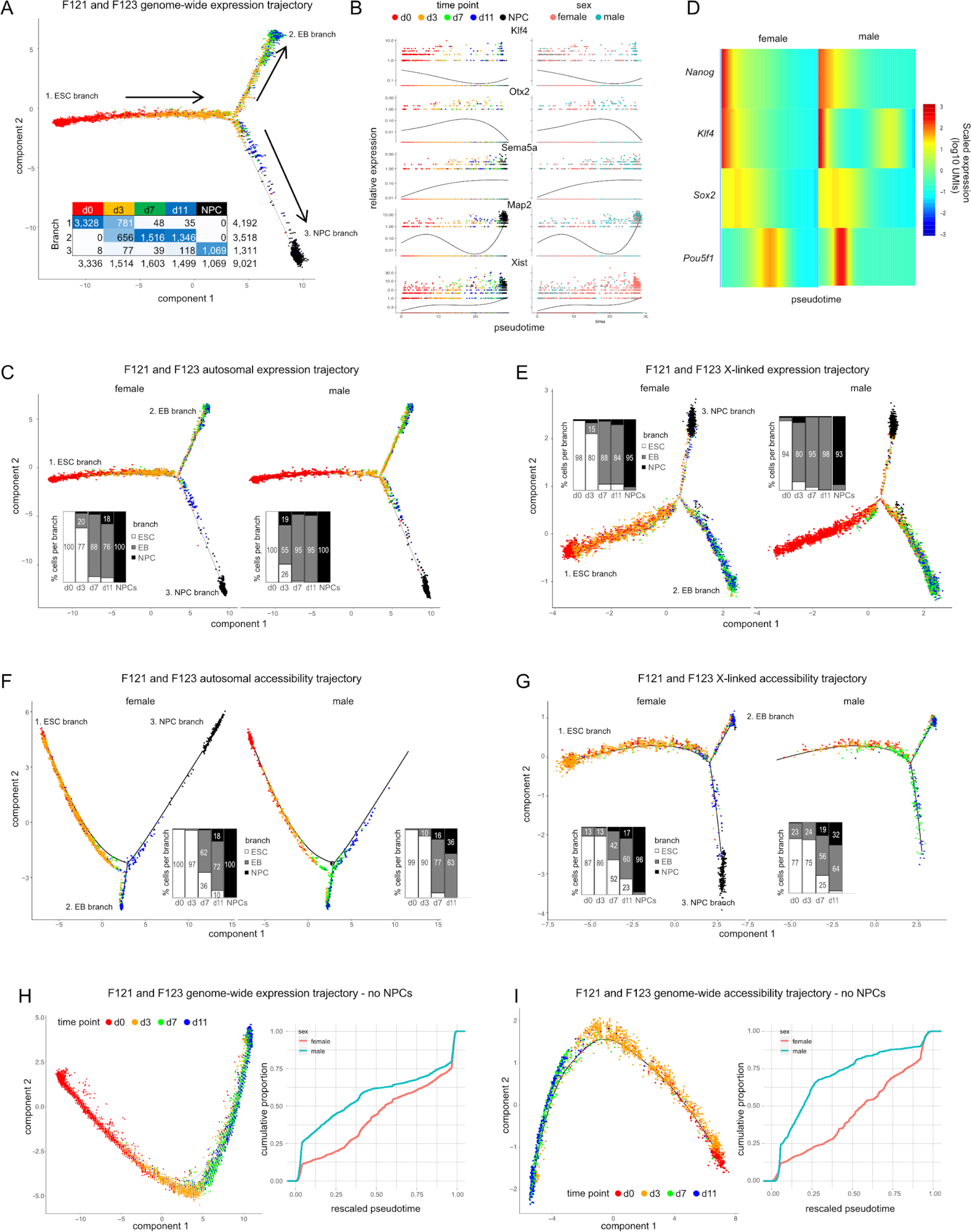
Female cells show delayed onset of differentiation relative to male cells. **A)** Semi-supervised genome-wide trajectory analysis with Monocle2 DDRTree using 14,037 differentially expressed genes across time points with q-value <= 0.01 and showing expression in at least 10 cells. Arrows indicate the direction of development in pseudotime. Branch 1 is the root ESC branch; branch 2 represents the default embryoid body (EB) pathway; and branch 3 captures NPCs, which derive from d11 cells. Inset table gives cell counts per time point per trajectory branch and time point color for data points in the plot. **B)** Expression (log10 UMIs) of select genes with respect to pseudotime based on the trajectory in A. *Klf4* is a pluripotency factor; *Otx2* is transiently expressed during EB differentiation; *Sema5a* is upregulated during EB differentiation; *Map2* is a NPC-specific marker; *Xist* is upregulated in female cells to initiate XCI. Data points are cells colored by time point (shown above) on the left, and by sex (female pink, male, blue) on the right. **C)** As in A, but based on 13,532 autosomal genes and faceted by cell sex (F121 on the left, F123 on the right). Bar graphs show the proportion of F121 to F123 cells per branch of the trajectory per time point. Compared to male cells female d3, d7, and d11 cells are enriched along the ESC branch 1, while d3 cells are under-enriched along the EB and NPC branches (branches 2 and 3) indicating a delayed onset of differentiation. Percentages of less than 10% omitted. **D)** Heatmap of genes in aligned kinetic curves where columns correspond to the log-transformed UMIs at each point in the aligned trajectory. These values were further transformed into per-gene Z scores. Each row shows a pluripotency factor gene. **E)** As in C, but for 361 X-linked genes. **F)** As in C, but based on chromatin accessibility data from sci-ATAC-seq for 2,919 of the top 25% most variable autosomal regions showing coverage in at least 10 cells and significantly differential across time points (q-value <= 10^−21^). **G)** As in F, but for 4,359 regions along chrX showing coverage in at least 10 cells and significantly differential across time points (q-value <= 0.01). **H)** As in A, but excluding NPCs. On the right are plots of cumulative distributions of cells’ pseudotimes (rescaled between 0 and 1) along the trajectory arc for female F121 and male F123 cells. **I)** As in H, but based on non-allelic chromatin accessibility.

We then investigated sex differences in pseudotime trajectories based on non-allelic autosomal gene expression patterns, which revealed differences in the developmental kinetics of female F121 and male F123 ESCs (Fig. 5C). Compared to male cells, female cells at d3, d7, and d11 are significantly enriched along the ESC branch, with 77% of female d3 cells still in that branch, as compared to 26% of male d3 cells. On the other hand, female cells at d3 are depleted along the EB branch where they represent only 20% of cells, while d3 male cells represent 55% of cells in that branch. Two genes encoding pluripotency factors (*Klf4, Pou5f1*) are exemplary of this delay in that they are expressed later in pseudotime in female versus male cells (Fig. 5D). An even more pronounced delay in female cell development is evident in each branch, especially between d0 and d3, when examining pseudotime trajectories based on X-linked gene expression, suggesting that the onset of XCI is a major contributor to the strong developmental delay (Fig. 5E). These sex differences were also observed in X-linked and autosomal chromatin accessibility patterns, providing further support for this early developmental sexual dimorphism (Fig. 5F, G). Although we cannot entirely rule out the possibility that this difference in early embryonic development between the male and female cells is due to differences in the developmental timing of these particular cell lines, it is consistent with previous studies that have shown a delay in the developmental program of female XX cells versus X0 or XY cells during early differentiation (Schulz et al. 2014)

Excluding NPCs from genome-wide non-allelic expression and accessibility analysis resulted in trajectories without branch points representing a smooth passage of development along the default pathway from ESCs to EBs (Fig. 5H, I). These latter trajectories also demonstrate the developmental delay experienced by female cells compared to male cells (Fig. 5H, I). To confirm results observed from the trajectories obtained from the DDRTree algorithm, we used an entirely unsupervised approach by conducting topic modeling on the F121 and F123 sci-RNA-seq and sci-ATAC-seq count matrices to determine common patterns of expression and accessibility across subsets of cells (see Methods). Projections of the resulting topic matrices into 2D space show that the topics successfully recapitulate the developmental time course, both by PCA and UMAP (Extended Data Fig. 6G, J). Segregating the cells by sex confirms that early in EB differentiation female cells – d3 cells in particular – are delayed in their development relative to male cells (Extended Data Fig. 6K–R).

Next, we generated allelic count matrices to assess the dynamics of gene expression and chromatin accessibility changes during female F121 and male F123 differentiation. We obtained 2,696 cells with at least 160 UMIs per allele by sci-RNA-seq (Extended Data Fig. 7A, B and Supplementary Table 3) and 1,900 cells with at least 500 UMIs by sci-ATAC-seq (Extended Data Fig. 7E, F and Supplementary Table 4). Distributions of allelic expression and accessibility were comparable for autosomes at all timepoints for both F121 and F123 cells, while X-linked distributions showed differences among developmental stages in female F121 cells. As expected, we also observed complete silencing of the 129 chrX in F121 NPSs, and absence of the *cast* allele in male F123 cells (Extended Data Fig. 7C, D, G, H). Heatmaps of allelic gene expression for the 25% most variability expressed genes on chr1 and chrX confirm these trends (Extended Data Fig. 7I, J). Trajectories based on genome-wide allelic expression and accessibility data (allelic counts concatenated together) in F121 cells are similar to those obtained for non-allelic data (Fig. 6A, B).

**Figure 6.**
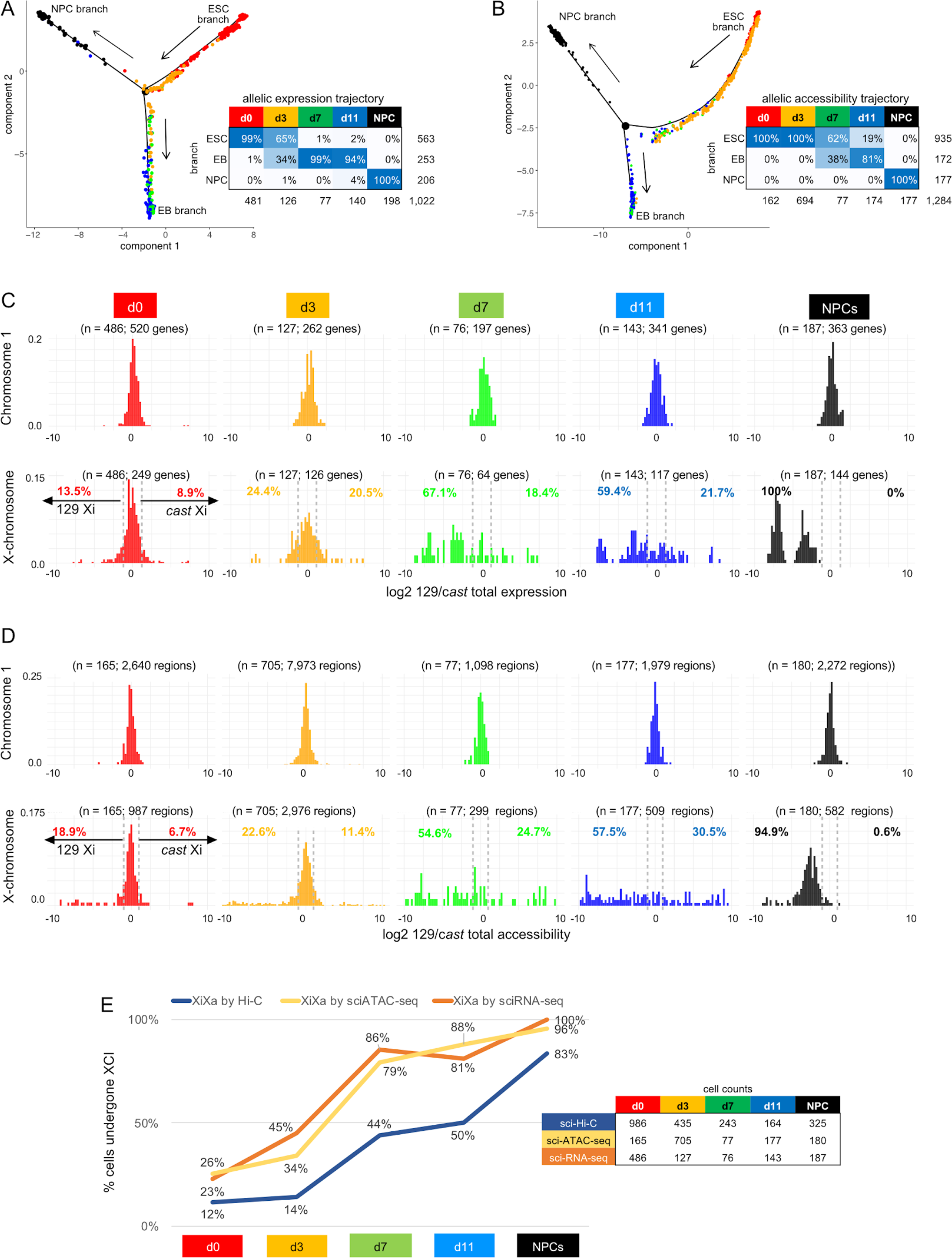
Allelic expression and accessibility distributions show the onset of XCI-induced gene silencing. **A)** Semi-supervised genome-wide trajectory analysis based on allelic expression data from sci-RNA-seq data with Monocle2 DDRTree using 4,230 differentially expressed alleles across time points (q-value <= 0.01) and expression in at least 10 cells with a minimum total allelic expression of 160 UMI. Arrows indicate the direction of development in pseudotime. Branches are labeled according to developmental state: ESC branch; EB branch, and NPC branch. Tables below trajectories give cell counts and percentages per time point per trajectory branch and time point color for data points in the plot. **B)** As in A, but based on allelic accessibility data from sci-ATAC-seq for 17,455 of the top 50% most variable regions across the genome showing coverage in at least 10 cells and significantly differential across time points (q-value <= 0.01). **C)** Distributions of differential expression from sci-RNA-seq between each homolog (log2 ratio of total 129 and *cast* expression) in female F121 cells at each time point (columns) for chr1 (top row) and chrX (bottom row) using genes expressed in at least 5 cells per group and cells with a total expression of at least 10 UMIs of expression from both alleles per chromosome. A 10% false positive rate (FPR) threshold (absolute log2 ratio >= 1.2) based on the chr1 distributions was used to classify cells that had undergone XCI based on the log2 ratios of their total allelic expression. Cells with either their 129 or *cast* allele having undergone XCI would have an allelic ratio skewed to the left and right, respectively, as indicated on the d0 plot. The percentage of cells exhibiting 129 or *cast* XCI at each time point is shown at the top left and right, respectively, of each chrX distribution. See Supplementary Table 2. **D)** As in C, but based on allelic chromatin accessibility data from sci-ATAC-seq. **E)** Plots of the proportion (%) of F121 cells with a silenced chrX (light orange line) as determined by sci-RNA-seq, with a chrX with low accessibility (yellow line) as determined by sci-ATAC-seq, and with an Xi bipartite structure (blue line) as determined by sci-Hi-C (see A and B and Fig. 3D, G).

Comparisons between the distributions of differential allelic expression and chromatin accessibility obtained by sci-RNA-seq and sci-ATAC-seq for chr1 and chrX (log2 ratio of total expression between parental alleles) show a switch from biallelic to monoallelic expression during differentiation for chrX, while chr1 shows biallelic expression (Fig. 6C, D). While autosomal distributions – such as those for chr1 – hug the zero value, consistent with biallelic expression, the distributions based on chrX allelic expression and accessibility spread away from zero during the course of differentiation, indicating an increasing proportion of cells with mono-allelic X-expression and accessibility due to the onset of XCI. Using a 10% FPR based on the chr1 distributions, one can estimate the proportion of cells with a silenced chrX as those with expression and accessibility skewed beyond the threshold in one direction or the other (129 or *cast*). As expected, this analysis shows that NPCs have XCI fully skewed towards the 129 allele in F121 cells. Plotting the proportion of cells with a silent chrX as a function of the time points highlights the onset of XCI, which shows a steep increase between d0 and d7 when 79-86% of cells have a silent chrX, followed by a slower increase to 96-100% of cells with a silent chrX in NPCs by sci-RNA-seq and sciATAC-seq (Fig. 6E and Supplementary Table 2). These proportions are robust to coverage level and cell number, with similar proportions at each time point observed using different sets of cells based on different total expression quantiles (Extended Data Fig. 7K – N). Importantly, comparison with the sci-Hi-C data shows a delay in the appearance of the Xi bipartite structure relative to silencing (Fig. 6E and Supplementary Table 2).

In summary, both the single-cell gene expression and chromatin accessibility data recapitulate the developmental trajectory of mouse ES cells to EB cells and NPCs. Our results also reveal that female cells lag behind their male counterparts, particularly early in the differentiation process, which is consistent with previous results (Schulz et al. 2014) and may reflect the establishment of XCI, but could also be due to inherent differences between the cell lines used. Allelic and non-allelic single-cell RNA-seq and ATAC-seq data produced similar trajectories that show the progressive onset of X-linked gene silencing in the course of differentiation. The proportion of cells with a silenced Xi increases in parallel to the proportion of cells exhibiting the Xi-specific 3D structure, but there is a lag in the appearance of the Xi structure.

### 5. Unsupervised multimodal alignment of single-cell expression, accessibility and contact data

Despite timing differences in the onset of silencing of X expression and of reduced chromatin accessibility versus the appearance of the Xi-specific structure, our single-cell measurements of distinct modalities captured a certain similarity in the dynamics of ESC differentiation and XCI features. Encouraged by these findings, we investigated the single-cell relationship between these modalities to better understand changes in chromatin structure and gene regulation during differentiation. For this analysis, we focused on F121 cells, which have a complete set of sci-Hi-C, sci-RNA-seq and sci-ATAC-seq data. We first aligned cells based on datasets obtained by sci-RNA-seq and sci-ATAC-seq separately from aliquots of the same batch of cells during differentiation. Considering that both experimental approaches appear to recapitulate the developmental trajectory and male-to-female difference (Fig. 5), we developed a method to align cells from each approach to, in effect, obtain an *in silico* co-assay. To align the cells, we employed the Maximum Mean Discrepancy Manifold Alignment (MMD-MA) algorithm (J. Liu et al. 2019). MMD-MA jointly embeds cells measured in different ways into a shared latent space to achieve an *in silico* co-assay without any supervision or prior knowledge as to the cell states (see Methods).

We successfully aligned cells based on non-allelic expression and chromatin accessibility patterns from F121 cells, as described in a separate publication (Singh et al. 2020). Here, we investigated whether we could extend our alignments between sci-RNA-seq and sci-ATAC-seq data to allelic data, and also to sci-Hi-C data. As we did with the non-allelic count matrices, we used topic modeling to extract the most prevalent patterns of F121 allelic expression and chromatin accessibility in order to denoise the data and thereby facilitate alignment (Extended Data Fig. 8A, B, D, E and see Methods). Similar to what was seen for the non-allelic topic matrices (Extended Data Fig. 6G–J), the allelic topics appear to capture the developmental trajectory of the cells. Furthermore, projections of the results of MMD-MA of F121 cells show that the algorithm was able to align the cells in a shared space, while retaining the cells’ inherent spatial separation with respect to time point (Fig. 7A). Even though the alignment was entirely unsupervised and did not utilize the time point information in performing the alignment, cells from similar time points clustered together in the new shared space. We evaluated alignment performance using the time-point labels as orthogonal information to calculate an AUC score (see Methods; Supplementary Table 5). The final mean AUC score for all cells in the datasets was 0.907, which is better than that obtained using non-allelic data (0.875).

**Figure 7.**
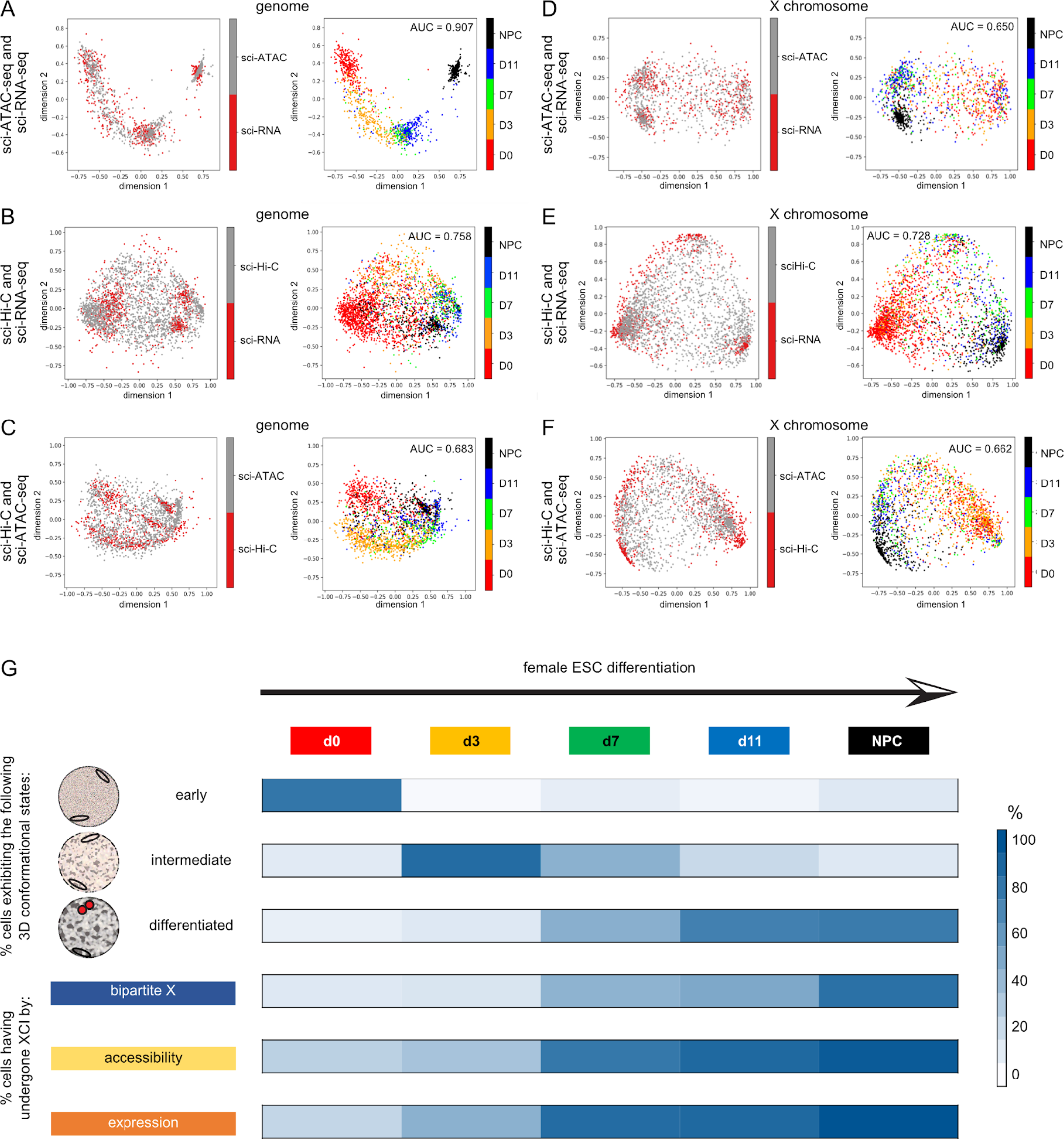
Alignment of allelic multimodal data during time course of ESC differentiation. **A)** Embedding of allelic sci-RNA-seq and sci-ATAC-seq topic matrices (see Extended Data Fig. 9A, B, D, E) learned by MMD-MA, projected in 2D via PCA. In the left-hand panel, each data point represents a cell colored by assay type. In the right-hand panel, each data point represents a cell colored by time point and the mean curve (AUC) score is shown (Supplementary Table 5; see Methods). **B)** Embedding of allelic sci-RNA-seq and sci-Hi-C topic matrices (see Extended Data Fig. 9A, C, D, F) learned by MMD-MA. Projection and panels as in A. **C)** Embedding of allelic sci-ATAC-seq and sci-Hi-C topic matrices (see Extended Data Fig. 9B, C, E, F) learned by MMD-MA. Projection and panels as in A. **D)** As in A, but only using chrX data. **E)** As in B, but only using chrX data. **F)** As in C, but only using chrX data. G) Schematic representation of the onset of structural changes throughout the genome and the X chromosome during female ESC differentiation. The proportions of cells with a specific feature are shown for each time point (labeled at top) by shades of blue corresponding to % of cells with that feature (Fig. 4A-C, 6E). Specific features are listed at left from top to bottom. Three types of genome-wide nuclear configuration detected by sciHi-C are schematized to reflect fewer long-range contacts in ESCs, partially condensed chromatin in the intermediate stage, and condensed chromatin in differentiated cells. The X chromosomes (ovals) appear to change structure at the same time as the rest of the genome, and their structure remains similar to that of other chromosomes in ESCs and at the intermediate stage. At the differentiated stage the Xi condenses along with the rest of the genome, but very-long range contacts are further enhanced to form the highly condensed Xi bipartite structure (red). Loss of chromatin accessibility and silencing of genes on the Xi proceed ahead of the structural changes.

In contrast to single-cell gene expression and chromatin accessibility data, non-allelic sci-Hi-C data proved more challenging to align to either of those datasets. We wondered whether using allelic sci-Hi-C data might improve the results since the allelic CDPs capture the onset and progressive establishment of the distinctive Xi architecture during differentiation (Fig. 3). A matrix of allelic sci-Hi-C-based concatenated CDPs within logarithmically-increasing sized bins for both alleles of each chromosome for each cell was transformed into a topic matrix. Projections of the sci-Hi-C topics appear to show patterns of interaction particular to each time point (Extended Data Fig. 8C, F). Although a clear developmental trajectory for the alignment of allelic expression and interaction data was not as obvious as seen for the alignment of allelic expression and accessibility data, MMD-MA was nonetheless able to align F121 cells with an AUC of 0.758 (Fig. 7B). The alignment of allelic accessibility and interaction data produced an AUC of 0.683 using genome-wide data (Fig. 7C).

To examine to what extent the alignments were driven by changes in the structure of the X chromosome, we performed them using just allelic chrX data (Fig. 7E, F). This analysis yielded AUCs of 0.728 and 0.662 for alignment of interaction data with gene expression and accessibility, respectively, nearly as high as those obtained using genome-wide data. Surprisingly, the alignment of cells based on allelic expression and accessibility using just X-linked data produced a much poorer alignment (AUC = 0.65) than that achieved using genome-scale data (AUC = 0.907) (Fig. 7D). Conversely, although the alignment of cells using allelic expression and accessibility data from all autosomes or chr1 alone produced results similar to that seen using genome-wide data (AUC = 0.911 and 0.848 respectively), alignment of cells based on autosomal or chr1 allelic expression and interaction data was relatively poor (AUC = 0.684 and 0.648 respectively) (Extended Data Fig. 8G – K). This suggests that the alignment of cells by allelic expression and 3D structure data is largely driven by the structural changes that occur to the future Xi during the course of differentiation due to XCI. Indeed, using data from both chrX and chr1 results in reasonable alignments based both on allelic expression and accessibility (AUC = 0.863) as well as based on allelic expression and interaction data (AUC = 0.732) (Extended Data Fig. 8I, L).

In conclusion, we were able to align single-cell datasets measuring three distinct modalities – gene expression, chromatin accessibility, and 3D chromosome organization – using unsupervised MMD-MA. To our knowledge this is the first time an *in silico* co-assay has been used to align three disjoint single-cell datasets. This multimodal alignment greatly improved clustering of cells to identify single-cell relationships among gene expression, chromatin accessibility, and nuclear structure during differentiation. The sci-RNA-seq and sci-ATAC-seq data show strong alignments as expected since the methods evaluate gene expression and associated *cis* regulatory changes respectively. In contrast, alignments with sci-Hi-C data reveal a number of cells that do not align, which could be due in part to nuclear structure variability. However, this may also reflect the observed lag in structural changes versus gene expression and chromatin accessibility changes during differentiation, as well as differences in the nature of the developmental changes observed - progressive gene expression and chromatin accessibility changes versus abrupt nuclear structure changes.

## Discussion

Allelic single-cell profiling of differentiating mouse ESCs based on gene expression, chromatin accessibility, and 3D chromosome structure combined with novel computational approaches to integrate these datasets provides a comprehensive roadmap of the dynamics of nuclear structure during differentiation and XCI. A major conclusion is that whereas gene expression and chromatin accessibility progressively change during differentiation, structural changes in nuclei are more abrupt and can be classified into three distinct stages based on the distribution of mid-range and long-range contacts (Fig. 7G). The establishment of a condensed bipartite Xi in which very-long range contacts are highest proceeds in parallel to those in the rest of the genome. Thus, all chromosomes including the Xi undergo a stepwise condensation of heterochromatic regions. Our single cell analyses reveal distinct nuclear configurations, including an early stage (ESCs at d0), an intermediate stage (d3 and d7 of EB differentiation), and a differentiated stage (NPCs and d11 of EB differentiation). This helps define three stages of chromosome configuration characterized by specific ratios of long-range versus medium-range contacts. The significant increase in long-range contacts in cells at the intermediate stage versus the early stage indicates the early formation of condensed heterochromatic regions, and the further increase in long-range contacts in cells at the differentiated stage marks a second phase of heterochromatin condensation.

Studies of early mammalian development *in vivo* have shown that chromatin exists in a relaxed state after fertilization, followed by progressive formation of higher-order chromatin architecture during early development up to embryo implantation (Du et al. 2017; Collombet et al. 2020; Ke et al. 2017). These bulk and single-cell Hi-C based analyses are focused on stages between gametes and preimplantation embryos, which show the appearance of TADs and the progressive formation of A/B compartments, resulting in a plaid-like structure in Hi-C contact maps. Mouse ESCs are derived from the inner cell mass of later embryos after implantation, and thus can serve as a model of subsequent developmental events. ESCs are known to be generally transcriptionally active, suggesting a dispersed chromatin structure correlated to pluripotency that becomes remodeled during differentiation and progressive gene silencing (Efroni et al. 2008). This is supported by a study based on electron spectroscopic imaging, which showed globally dispersed chromatin in cells from the inner cell mass and in ESCs from embryonic stage E3.5, in contrast to compacted chromatin located near the nuclear envelope in epiblasts and epiblast-like stem cells at embryonic stage E5.5 (Ahmed et al. 2010). A similar study based on the differentiation of ESCs to NPCs confirmed this transformation in chromatin organization (Hiratani et al. 2010). A pluripotency-specific genome organization has also been recapitulated during reprogramming from a differentiated state (Apostolou et al. 2013; Denholtz et al. 2013). Furthermore, consistent with our results, bulk studies of mouse ESC differentiation show evidence of A/B compartments, which become stronger during differentiation in NPCs and are further enhanced in cortical neurons and brain (Bonev et al. 2017; Miura et al. 2019). This global heterochromatin formation and condensation is correlated to changes from early to late replication timing and subnuclear positioning during differentiation (Miura et al. 2019; Hiratani et al. 2010).

We find that the Xi-specific bipartite structure appears simultaneously with changes in contact distribution for other chromosomes. Thus, we observed three distinct stages of condensation of the Xi concurrent to those observed in the rest of the genome (Fig. 7G). The Xi-specific bipartite structure becomes apparent in differentiating female ESCs at d7 but is not prevalent at d3, suggesting that late-stage events such as macroH2A enrichment, SMCHD1-mediated DNA methylation, and late DNA replication may assist the assembly of the Xi bipartite structure (Gendrel et al. 2012; Wang et al. 2018; Gdula et al. 2019; Mermoud et al. 1999; Miura et al. 2019). In contrast, *Xist* coating very quickly initiates silencing via early histone modifications (Żylicz et al. 2019). Bulk studies of mESCs have suggested the existence of an intermediate stage of Xi condensation via S1 and S2 compartments, which would then fuse in the presence of SMCHD1 (Wang et al. 2018). Whether these S1/S2 compartments potentially represent the intermediate stage we detected is unclear. Interestingly, SMCHD1 is recruited to the Xi at d7 with only ∼20% of cells showing Xi enrichment, but then increases rapidly to ∼80% at d9 (Gendrel et al. 2012). Our trajectory analyses show that cells at d11 and NPCs cluster at the differentiated branch based on their CDPs. Additional changes in heterochromatin condensation in specific cell types in different lineages remain to be characterized.

Based on alignment of datasets and on trajectories we show that the timing of initiation of the Xi-specific structure is variable among cells and lags behind *Xist* RNA coating and silencing of the Xi. Indeed, more than 80% of ES_Tsix-stop cells already have an *Xist* RNA cloud at d3, but only 34% have acquired an Xi structure (this study) (Monkhorst et al. 2008). The higher proportion of cells showing an Xi marked by *Xist* using microscopy compared to those with a detectable bipartite structure by sci-Hi-C could be due to methodological differences. Certainly *Xist* upregulation and cis-coating on the Xi would be expected to be an early event. However, as discussed above, the establishment of a full bipartite structure requires late events to assemble the heterochromatic structure (Nora and Heard 2010; Wang et al. 2018; Gendrel et al. 2012; Mermoud et al. 1999; Miura et al. 2019). The presence of a few cells scored as positive for a bipartite X at d0 could be due to the stringent threshold chosen, but could also reflect the presence of a bipartite X in a few undifferentiated cells. There is evidence that the X chromosomes may switch between two different states prior to XCI in a proportion of mouse ESCs, as a way to initiate random XCI upon differentiation (Mlynarczyk-Evans et al. 2006).

While *Xist* and H3K27me3 remain associated with the mouse Xi through the cell cycle, our results show a rapid loss of the Xi bipartite structure at mitosis (Duthie et al. 1999). The Xi bipartite structure reappears quickly upon exit from mitosis, suggesting a possible ‘bookkeeping’ mechanism of the Xi bipartite structure, which could be mediated by CTCF binding at *Dxz4*. CTCF bookmarking sites in ES cells are abundant and generally involved in rapid gene reactivation after replication and mitosis (Owens et al. 2019). In contrast, somatic cells have few such sites, which are involved in maintaining chromatin structure, for example, specific loops at imprinted genes (Y. Zhang et al. 2019); Cai et al., 2018 Nature; Burke et al., EMBO J. 2005).

By single-cell RNA-seq and ATAC-seq we clearly detected a delay in the differentiation of female versus male cells, a finding consistent with a differentiation block in female cells due to the presence of two active X chromosomes (Schulz et al. 2014). This delay shows cell-to-cell heterogeneity and is inversely correlated with upregulation of *Xist* and initiation of XCI, as evidenced by the detection of two groups of female cells at d3, including one group with high *Xist* expression and similar transcriptome and chromatin accessibility kinetics as male cells, and a second group with low *Xist* expression and slower expression kinetics. The female-specific delay almost disappears at d7 and d11, suggesting that dosage compensation by XCI of genes related to differentiation may release this delay. Interestingly, subtle differences in X-linked and autosomal expression profiles were observed between male and female cells at d7, d11, and NPCs based on UMAP-based clustering analyses, suggesting that sex differences persist at later differentiation time points due to genes that escape XCI in females and/or the presence of Y-linked genes in males. However, these findings should be confirmed in additional cell lines.

In this study, cells were aligned at each time point of differentiation and along developmental trajectories characterized by specific gene expression, chromatin accessibility, and 3D chromosome structure, effectively providing in silico co-assays. We found that trajectories that reflect differentiation branches could be built using allelic, but not non-allelic, sci-Hi-C data. Trajectories were obtained not only for the X chromosomes, but also for the whole genome and for autosomes only, suggesting that inherent structural differences between homologs exist, possibly due to imprinted genes or other mono-allelicallly expressed genes (Chess 2016; X. Deng et al. 2015). We also cannot exclude indirect effects of the heterochromatic Xi on the 3D structure of the rest of the genome. Our alignment approach will help understand the relationships between the 3D nuclear structure, chromatin features, and the transcriptome in various cell types and during development.

## Methods

### Cell lines

Patski cells are fibroblasts derived from 18dpc embryonic kidney from a cross between a *Mus musculus* strain C57BL/6J (B6) female with an *Hprt*^*BM3*^ mutation and a *Mus spretus* (*spret*) male (Figure 1). Cells have been selected in HAT (hypoxanthine-aminopterin-thymidine) media such that the B6 chrX is always inactive, as verified in previous studies (Lingenfelter et al. 1998; G. Bonora et al. 2018). Patski cells were cultured in Dulbecco’s Modified Eagle Medium (DMEM) with 10% FBS and 1% penicillin/streptomycin.

Mouse embryonic stem cells (mESCs) F121-6 (female; F121 hereafter) and F123 (male) derived from a cross between *Mus musculus* strain 129/SV-Jae (129) and *Mus castaneus* (*cast*) were obtained from J. Gribnau. A second female F1 hybrid ESC line, ES_Tsix-stop, was derived from the same cross between 129 and *cast*, followed by the introduction of a transcriptional stop in the *Tsix* gene on the 129 chrX (Figure 1) (Luikenhuis, Wutz, and Jaenisch 2001). In cells differentiated from ES_Tsix-stop the Xi is always from 129. All mESCs were cultured and expanded on MEF feeders following the standard operating procedure designed for F123 (https://www.4dnucleome.org/cell-lines.html), except for the addition of 1% penicillin/streptomycin.

Differentiation of mESCs into EBs was done using a standard LIF-withdrawal protocol. In brief, mESCs were grown on gelatin-coated plates for two passages to remove MEF feeders. Feeder-free mESCs were harvested and aliquots saved as day 0 (d0). The remaining cells were cultured in regular medium (10% FBS and 1% penicillin/streptomycin in DMEM with high glucose) in non-adherent petri-dishes for 11 days of EB differentiation. EBs were collected at days 3, 7, and 11 (d3, d7, and d11) and treated with accutase to prepare single-cell suspensions (Fig. 1). An aliquot of d11 EBs was used to derive NPCs using a published protocol with slight modifications (Xu et al. 2005). In brief, accutase-treated d11 EBs were plated onto adhesive tissue culture dishes precoated with 1% gelatin in regular medium. Selection of NPCs was initiated 24h later by replacing the medium with serum-free selection medium consisting of 1:1 DMEM/F12:neurobasal medium plus 1xN2 supplement (GIBCO, 50x) for ∼5 days. After selection, cells were harvested and grown for 5-6 passages in NPC expansion medium (selection medium with the addition of 2% B27 [GIBCO], 20 ng/ml bFGF [Sigma], and 20 ng/ml EGF [Sigma]) before collection for single-cell analysis. NPCs were identified by staining with an Alexa Fluor 488 conjugated antibody for nestin (Millipore Cat. # MAB353A4; Extended Data Fig. 9). A clonal F121 NPC line was derived from the bulk culture by single-cell-patch picking. This NPC clone was found to carry an Xi from 129 in all cells as shown by SNP analysis of sci-RNA-seq. Establishment of XCI in female ES cells (F121 and ES_Tsix-stop) and control male cells F123 collected from d0 to d11 and from NPCs was tested by *Xist* RNA FISH and verified in NPCs by H3K27me3 immunofluorescence to detect the Xi (Extended Data Fig. 9). *Xist* RNA-FISH was done using a 10kb *Xist* cDNA plasmid (pXho, which contains most of exon 1 of *Xist*) as described (Yang et al. 2015). Immunostaining of paraformaldehyde-fixed nuclei using an H3K27me3 antibody (Millipore Cat. # 07-449) was done in female NPCs as described (Yang et al. 2015). About 100 or more nuclei were scored for the presence/absence of *Xist* RNA clouds or for the presence of an enriched H3K27me3 cluster (Extended Data Fig. 9). EB differentiation of mESCs (F123, F121, and ES_Tsix-stop) was repeated to generate replicates by collecting cells at d0 and d7. In addition, d7 EBs were plated back to adhesive cell culture dishes to allow EB attachment and outgrowth of differentiated fibroblasts, which were collected at d20.

### sci-Hi-C

Sci-Hi-C was done according to published protocols (Ramani et al. 2017, 2020). To control for aberrant cell aggregation human cells were mixed with the mouse cells. Sci-Hi-C libraries were processed using a publicly available pipeline (Ramani et al. 2017, 2020), which was adapted to process allelically segregated reads. Reads were aligned to an N-masked C57BL6/J reference genome (mm10) where every SNP locus (129 or *cast* for the F123, F121, and ES_Tsix-stop cell lines; B6 or *spret* for the Patski cell line) was substituted with an N to reduce mapping bias. The pipeline resulted in lists of contact pairs for all cells with at least 1,000 uniquely mapped contact pairs, a *cis:trans* ratio ≥1 and ≥95% of reads mapping to either the mouse or human genome (Supplementary Table 1). Reads from human cells were used to estimate the doublet rate and to filter cells with high alignment rates to both the mouse and human reference genomes. Contact matrices for the mouse genome were generated from the valid pairs by summing counts within bins of a specified size (e.g. 500kb). To obtain allelic information, each valid contact pair was segregated to its parental allele of origin if either end of the read pair contained at least one SNP particular to one of the parental strains. SNPs were based on Sanger data for mouse strain 129 and mouse species *M. spretus* and *M. castaneus*, compared to the mm10 B6 reference assembly (Keane et al. 2011; Lilue et al. 2018). Read pairs containing no SNPs or containing SNPs belonging to both parental strains could not be unambiguously assigned to a single allele and were discarded.

Contact decay profiles (CDPs) for each cell were generated by summing the biallelic intrachromosomal contact counts with respect to contact distance within exponentially increasing contact distance ranges, as described (Nagano et al. 2017) (Extended Data Fig. 2A, B). Specifically, bin boundaries corresponded to the series of distances 2^x^ to 2^(x+1)^ for x within the range [10; 27.750] incrementing by a step size of 0.125. Cells were grouped into four clusters by k-means clustering using the Spearman correlation distance between the CDPs within the 50kb to 8Mb range for each cell. The four clusters correspond to cells in different stages of the cell cycle progressing from mitosis into different stages of interphase.

In differentiated cells (e.g. Patski and NPCs) in which XCI is complete, the Xi shows a distinct CDP profile compared to that of the Xa and autosomal alleles. Specifically, the Xi CDP profile shows a lower proportion of mid-range contacts (85 kb - 1.1 Mb) and a higher proportion of very long-range contacts (6.5 Mb - 87 Mb). This difference was made even more evident when noise due to sparsity was reduced by further aggregating the contact counts into non-overlapping sets of 10 logarithmic bins for each homolog of each pair of chromosomes of each cell. A Spearman correlation between the resulting rebinned allelic CDPs for each homolog was calculated for each chromosome of every cell with sufficient coverage, i.e., at least 100 intrachromosomal contacts from each allele in Patski cells and at least 50 intrachromosomal contacts from each allele in the other cell types analyzed together. A lower Spearman correlation coefficient between the rebinned CDPs of the Xi and Xa is indicative of the Xi having assumed a 3D conformation of a cell that had undergone XCI. A threshold value (ρXi ^CDP^) was chosen based on a 10% false positive rate (FPR) using the distribution of Spearman correlation coefficients between the rebinned allelic CDPs of the chr1 homologs, which would be expected to show a high CDP correlation. Cells with a Spearman correlation between the rebinned allelic CDPs of the chrX homologs below ρXi^CDP^ were deemed to have one chrX homolog that had assumed the Xi 3D structure, but the identity of this homolog was unknown in the case of cells with random XCI.

To determine which chrX homolog had acquired a 3D structure characteristic of the Xi in cells such as F121 cells, where XCI is only partially skewed during differentiation, the difference between the total long-range (6.5Mb - 87Mb) and mid-range (85kb - 1.1Mb) contact counts was assessed for each homolog of each cell, referred to hereafter as the long-range to mid-range difference (LMD). The difference between the resulting allelic LMDs for each homolog was calculated for each chromosome of every cell with sufficient coverage, in the same fashion as the Spearman correlation analysis. A large LMD difference between the rebinned CDPs of chrX homologs A and B, say, is indicative of the fact that homolog A has assumed the Xi 3D structure. A threshold value (δ Xi^LMD^) was chosen based on a 10% FPR using the distribution of LMD differences between the rebinned allelic CDPs of the chr1 homologs, which would be expected to show little difference in their LMDs. Cells with an absolute LMD difference between the rebinned allelic CDPs of the chrX homologs greater than δ Xi^LMD^ were deemed to have an chrX homolog that had assumed the Xi 3D structure and the homolog with the higher LMD was classified as the Xi. This LMD analysis in cells with skewed XCI (Patski cells, cloned F121 NPCs, and differentiated ES_Tsix-stop cells) successfully identified the default chrX homolog to be the Xi, assuring the accuracy and efficiency of this method in classification of the Xi based on 3D structures.

To count the number of read pairs spanning each locus along a chromosome across all contact length scales out to a set distance, we used a “contact score” – an adaptation to the “coverage score” metric used in (Galupa and Heard 2018; Giancarlo Bonora and Disteche 2017). Briefly, the contact score for each bin at a particular resolution was calculated as the average number of interactions within bins spanning this central bin of interest. This score calculation can be visualized as sliding a V-shaped region (with the arms of the V extending out 20Mb) along the diagonal of the contact map and calculating the mean interaction counts (sum of counts/number of bins) within the region. We excluded bins within the first and last 10Mb of the chromosome in order to avoid edge effects. In addition, a pseudocount of 1 was added to all bins. The scores were normalized by calculating log2(coverage score/chromosomal mean), and a five-window, degree two polynomial Savitzky-Golay filter was applied to the resulting vector to smooth the signal. For the purposes of smoothing, bins without values were spline interpolated but replaced with NaNs after smoothing.

### sci-RNA-seq

Sci-RNA-seq was performed on nuclei isolated from non-fixed frozen cell aliquots using a published protocol (Cao et al. 2017, 2019). Sci-RNA-seq libraries were processed using a pipeline by the Trapnell lab (Cao et al. 2017), adapted to handle allelically segregated reads, including aligning reads to an N-masked B6 reference genome (mm10), as described above. The pipeline produces CellDataSet (CDS) files that contain genes-by-cell count matrices of unique molecular identifiers (UMIs) per gene, which were used for all downstream analysis. Genes with coverage along one allele in at least 10 cells and cells with at least 10 UMIs per each chr1 and chrX allele were included for downstream analysis. Mapped single-end reads were segregated to their allele of origin if the read contained at least one SNP particular to exactly one of the parental strains. Reads containing no SNPs or containing SNPs belonging to both parental strains were discarded. Genes expressed from one allele in at least 10 cells and cells with at least 10 UMIs in chr1 and in chrX were included (Supplementary Table 3).

To assess the XCI status of each cell, the total allelic expression (TAE; i.e. total UMI counts) for each chrX homolog was calculated, and the log2 ratio between the total expression between the two homologs was determined. A large difference in the TAE between chrX homologs A and B, say lower for B, is indicative of the fact that homolog B has been silenced. Using a similar approach to that used to identify significantly different allelic CDPs, a differential expression threshold (δ Xi^*TAE*^) was chosen based on a 10% FPR using the distribution of TAE log2 ratios between the rebinned allelic CDPs of the chr1 homologs, which would be expected to show biallelic expression. Cells with an absolute TAE log2 ratio between their chrX homologs of greater than δ Xi^*TAE*^ were deemed to have an chrX homolog that had undergone XCI, and the homolog with the higher TAE was classified as the Xa.

### sci-ATAC-seq

Sci-ATAC-seq was done using a published protocol (Cusanovich et al. 2018). Sci-ATAC-seq libraries were processed using a pipeline created by the Trapnell lab (https://github.com/cole-trapnell-lab/ATAC-Pipeline-Dev), which we adapted to handle allelically segregated reads, including aligning reads to an N-masked B6 reference genome (mm10), as described above for sci-Hi-C. Similar to the sci-RNA-seq pipeline, the sci-ATAC-seq pipeline produces CDS files that contain count matrices of UMIs per accessible region. These regions are defined as peaks of accessibility called using MACS2 on the aggregated sci-ATAC-seq data. Accessible sites with coverage in at least 10 cells were included. The CDS files were used for all downstream analysis. Aligned paired-end reads were segregated to their allele of origin as described above. Accessible regions with coverage along one allele in at least 10 cells and cells with at least 10 UMIs per each chr1 and chrX allele were included (Supplementary Table 4).

To assess the XCI status of each cell, the total allelic accessibility (TAA; i.e. total UMI counts) for each chrX homolog was calculated, and the log2 ratio between the total accessibility between the two homologs was determined. A large log2 ratio in the TAA between chrX homologs A and B, say greater for A, is indicative of the fact that homolog B has been silenced. As was done for the TAE (described above), a differential accessibility threshold (δ Xi^*TAA*^) was chosen based on a 10% FPR using the distribution of TAA log2 ratios between the rebinned allelic CDPs of the chr1 homologs, which would be expected to show equal levels of accessibility. Cells with an absolute TAA log2 ratio between their chrX homologs of greater than δ Xi^*TAA*^ were deemed to have an chrX homolog that had undergone XCI, and the homolog with the higher TAA was classified as the Xa.

### Dimensionality reductions, topic modeling and trajectory analyses

Dimensionality reduction was performed using both PCA and Uniform Manifold Approximation and Projection (UMAP) using default parameters (McInnes et al. 2018). Trajectory analyses were generated by Monocle2 (Qiu et al. 2017) using the DDRtree algorithm (Mao et al. 2015) in semi-supervised mode using differentially expressed genes across the time points.

Topic modeling was performed on binarized feature-by-cell count matrices using cisTopic (Bravo González-Blas et al. 2019)). CisTopic employs a Latent Dirichlet Allocation (LDA) topic modeling algorithm, which decomposes a gene-by-cell count matrix into a topics-by-cell matrix and a cells-by-region matrix, with the topics capturing the most prevalent patterns of features in cells thereby enhancing cell type-specific signals in the data. The runCGSModels() was applied to the binarized count matrices with default parameters (burnin = 250, iterations = 500). The model with the maximum log-likelihood was always selected as the number of topics.

### Integration of datasets by MMD-MA

To align the F121 cells, which have sci-RNA, sci-ATAC and sci-Hi-C data at five time points, we employed the Maximum Mean Discrepancy Manifold Alignment (MMD-MA) algorithm (J. Liu et al. 2019; Singh et al. 2020). The raw data inputs were as follows:

- Non-allelic expression data: a gene-by-cell count matrix of sci-RNA-seq UMI tallies within genes (rows) of each cell (columns) was used as input. Genes expressed in at least 10 cells were included.
- Non-allelic accessibility data: a matrix of sci-ATAC-seq UMI counts within accessible regions (rows) of each cell (columns) was used as input. Accessible sites with coverage in at least 10 cells were included.
- Allelic expression data: a matrix of sci-RNA-seq UMI counts within the genes from each parental allele were concatenated together (rows) for each cell (columns). Genes expressed from a parental allele in at least 10 cells and cells with at least 10 UMIs in chr1 and chrX were included.
- Allelic accessibility data: a matrix of sci-ATAC-seq UMI counts within accessible regions from each parental allele concatenated together (rows) for each cell (columns). Accessible regions with coverage along a parental allele in at least 10 cells and cells with at least 10 UMIs per each chr1 and chrX allele were included.
- Allelic DNA interaction data: a matrix of sci-Hi-C-based concatenated contact decay counts within logarithmically-increasing sized bins for both parental alleles of each chromosome (rows) for each cell (columns). Only cells with at least 50 contacts in each allele from chr1 and chrX were included.

Prior to alignment, each of the aforementioned count matrices were decomposed into topic-by-cell and feature-by-topic matrices using cisTopic (Bravo González-Blas et al. 2019)). The topic-by-cell matrices served as input to the MMD-MA algorithm, as in (Singh et al. 2020). A key assumption of MMD-MA is that the data is sampled from the same underlying distribution, so before applying MMD-MA to our sci-RNA-seq and sci-ATAC-seq measurements, we first subsampled the cells from the five time-points (d0, d3, d7, d11, NPCs) to ensure that the fraction of cells in each time-point was consistent across the two datasets (Supplementary Table 5).

The MMD-MA algorithm aligns the single-cell data points from multiple domains by optimizing an objective function that reduces the overall similarity of the projected cell distributions, as measured by MMD. To select hyperparameters, including the number of dimensions of the shared latent space, we optimized the segregation of cells by their respective time points. We optimized the MMD-MA algorithm for the range of hyperparameters used in (Singh et al. 2020), including the size of the latent space in which the two datasets were aligned (Supplementary Table 5).

We evaluated the algorithm’s alignment performance using the time-point labels as orthogonal information to calculate the area under the curve (AUC) of a receiver operating characteristic curve (Supplementary Table 5). For each point in the aligned space, we calculate its Euclidean distance from other points, using them as scores. Next, we assign binary labels such that if a point belongs to the specific time-point cluster, it is assigned label ‘1’ else ‘0’. Hence, for a point in a time-point cluster, the AUC score calculation measures whether other points in the same cluster are closer than (hence ranked higher than) the points outside the cluster. This evaluation is done separately for each time point (relative to all other time points), and the four AUC scores are averaged. Therefore, in the learned latent space where both datasets have been aligned, the AUC analysis quantifies how well cells from the same time-point but measured in different modalities occur near one another. We split the data into training (70%) and validation sets (30%) and used the AUC score metric on the validation data to select the best performing hyperparameters.

## Data Availability

All data has been deposited into the 4DN data portal and will be available via the following URL: http://data.4dnucleome.org/bonora_sc-differentiation

## Code Availability

All code used in this project will be put into a publicly available repository upon publication of the manuscript.

## Acknowledgements

This study was supported by grants DK107979 (JS and WSN) and HG011586 (WSN, JS, and CMD) from the National Institutes of Health Common Fund 4D Nucleome, and by grants GM131745 (CMD) and GM1273727 (XD) from the National Institute of General Medical Sciences. JS is an Investigator of the Howard Hughes Medical Institute. We thank J. Gribnau (Erasmus University) for the mouse F1 hybrid ESC lines, F121-6, F123, and ES_Tsix-stop. We thank L. Saunders (University of Washington) for her assistance with the generation of sci-RNA-seq libraries, and J. Packer and H. Pliner (University of Washington) for assistance with the sci-RNA-seq and sci-ATAC-seq data processing pipelines.

