## Supplementary and Extended Data for "Single-cell landscape of nuclear configuration and gene expression during stem cell differentiation and X inactivation"

### Supplementary information and extended data files

### Supplementary Tables

***Supplementary Table 1. sci-Hi-C cell counts and non-allelic and allelic contact count summary statistics.***

| | | Non-allelic interaction summary statistics | | | | | |
|---|---|---|---|---|---|---|---|
| | | Genome-wide | | | X-chromosome | | |
| sample | cell counts | total contacts | mean contacts | median contacts | total contacts | mean contacts | median contacts |
| Patski | 4,493 | 39,289,629 | 8,745 | 5,167 | 2,153,390 | 479 | 280 |
| TsixStop_d0 | 342 | 1,770,062 | 5,176 | 4,491 | 85,204 | 249 | 223 |
| TsixStop_d3 | 839 | 2,878,080 | 3,430 | 2,785 | 134,247 | 160 | 128 |
| TsixStop_d7 | 822 | 4,748,683 | 5,777 | 4,218 | 232,140 | 283 | 199 |
| TsixStop_d11 | 471 | 3,517,515 | 7,468 | 3,392 | 176,193 | 374 | 153 |
| TsixStop_NPC | 2,535 | 16,481,251 | 6,501 | 4,903 | 834,206 | 329 | 246 |
| F121_d0 | 2,203 | 17,874,237 | 8,114 | 6,245 | 890,230 | 404 | 305 |
| F121_d3 | 894 | 9,688,584 | 10,837 | 6,297 | 578,606 | 647 | 367 |
| F121_d7 | 732 | 10,684,487 | 14,596 | 6,866 | 494,258 | 675 | 302 |
| F121_d11 | 608 | 7,266,390 | 11,951 | 6,064 | 314,722 | 518 | 260 |
| F121_NPC | 1,054 | 6,369,538 | 6,043 | 4,063 | 338,132 | 321 | 214 |
| F123_d0 | 1,670 | 14,520,800 | 8,695 | 6,668 | 421,215 | 252 | 190 |
| F123_NPC | 1,030 | 6,662,438 | 6,468 | 5,167 | 192,234 | 187 | 152 |
| replicate TsixStop_d0 | 795 | 8,777,503 | 11,041 | 6,339 | 388,210 | 488 | 262 |
| replicate TsixStop_d7 | 152 | 1,751,101 | 11,520 | 7,638 | 74,368 | 493 | 322 |
| replicate TsixStop_d20 | 404 | 5,348,989 | 13,240 | 3,951 | 257,677 | 638 | 189 |
| replicate F121_d0 | 1,479 | 8,469,522 | 5,727 | 3,707 | 455,156 | 308 | 200 |
| replicate F121_d7 | 89 | 2,279,421 | 25,611 | 7,739 | 111,292 | 1,250 | 382 |
| replicate F121_d20 | 301 | 3,144,889 | 10,448 | 4,643 | 158,900 | 530 | 227 |

| | | Allelic interaction summary statistics | | | | | | |
|---|---|---|---|---|---|---|---|---|
| | | Genome-wide | | | | X-chromosome | | |
| sample | cell counts | B6/129 contacts | spretus/Cast contacts | total allelic contacts | % allelic contacts | B6/129 contacts | spretus/Cast contacts | total allelic contacts |
| Patski | 4,493 | 15,141,925 | 11,144,006 | 26,285,931 | 67% | 714,755 | 575,179 | 1,289,934 |
| TsixStop_d0 | 342 | 560,583 | 528,075 | 1,088,658 | 62% | 17,984 | 18,977 | 36,961 |
| TsixStop_d3 | 839 | 907,602 | 855,558 | 1,763,160 | 61% | 27,939 | 30,936 | 58,875 |
| TsixStop_d7 | 822 | 1,478,343 | 1,397,457 | 2,875,800 | 61% | 48,383 | 56,053 | 104,436 |
| TsixStop_d11 | 471 | 1,096,305 | 1,028,002 | 2,124,307 | 60% | 36,138 | 42,144 | 78,282 |
| TsixStop_NPC | 2,535 | 5,192,202 | 4,853,045 | 10,045,247 | 61% | 185,676 | 182,927 | 368,603 |
| F121_d0 | 2,203 | 5,732,612 | 5,069,354 | 10,801,966 | 60% | 200,624 | 186,690 | 387,314 |
| F121_d3 | 894 | 2,875,116 | 2,576,480 | 5,451,596 | 56% | 121,324 | 114,033 | 235,357 |
| F121_d7 | 732 | 3,161,922 | 2,869,610 | 6,031,532 | 56% | 89,467 | 113,557 | 203,024 |
| F121_d11 | 608 | 2,148,784 | 1,957,360 | 4,106,144 | 57% | 55,715 | 72,183 | 127,898 |
| F121_NPC | 1,054 | 2,051,798 | 1,818,259 | 3,870,057 | 61% | 74,618 | 75,030 | 149,648 |
| F123_d0 | 1,670 | 4,757,858 | 4,115,468 | 8,873,326 | 61% | 175,901 | 12,429 | 188,330 |
| F123_NPC | 1,030 | 2,214,102 | 1,874,402 | 4,088,504 | 61% | 80,332 | 5,816 | 86,148 |
| replicate TsixStop_d0 | 795 | 2,629,500 | 2,428,991 | 5,058,491 | 58% | 76,029 | 85,089 | 161,118 |
| replicate TsixStop_d7 | 152 | 519,105 | 485,130 | 1,004,235 | 57% | 12,194 | 18,585 | 30,779 |
| replicate TsixStop_d20 | 404 | 1,572,477 | 1,506,863 | 3,079,340 | 58% | 48,830 | 60,170 | 109,000 |
| replicate F121_d0 | 1,479 | 2,597,383 | 2,309,070 | 4,906,453 | 58% | 98,774 | 89,158 | 187,932 |
| replicate F121_d7 | 89 | 688,096 | 621,063 | 1,309,159 | 57% | 21,981 | 24,050 | 46,031 |
| replicate F121_d20 | 301 | 943,181 | 849,736 | 1,792,917 | 57% | 34,696 | 32,453 | 67,149 |

Counts of cells with at least 1,000 uniquely mapped contact pairs, a *cis:trans* ratio ≥1 and ≥95% of reads mapping to either the mouse or human genome (see Methods) for each sci-Hi-C library, along with the total, mean, and median number of contact pairs per cell genome-wide and along chrX. The number of allelic contacts for each allele and in total is also provided for the genome (along the percentage of non-allelic contacts) and chrX.

***Supplementary Table 2. Counts and percentages of cells showing XCI by allelic sci-Hi-C, sci-RNA-seq, and sci-ATAC-seq.***

| dataset | XCI based on single cell interactions | | | | | | |
|---|---|---|---|---|---|---|---|
| | total cells | XCI cells | B6/129 Xi cells | Spret/Cast Xi cells | XCI % | B6/129 Xi % | Spret/Cast Xi % |
| F121_d0 | 986 | 116 | 59 | 57 | 12% | 6% | 6% |
| F121_d3 | 435 | 61 | 34 | 27 | 14% | 8% | 6% |
| F121_d7 | 243 | 107 | 68 | 39 | 44% | 28% | 16% |
| F121_d11 | 164 | 82 | 52 | 30 | 50% | 32% | 18% |
| F121_NPC | 325 | 271 | 271 | 0 | 83% | 83% | 0% |
| TsixStop_d0 | 89 | 11 | 8 | 3 | 12% | 9% | 3% |
| TsixStop_d3 | 86 | 30 | 29 | 1 | 35% | 34% | 1% |
| TsixStop_d7 | 159 | 103 | 100 | 3 | 65% | 63% | 2% |
| TsixStop_d11 | 83 | 60 | 59 | 1 | 72% | 71% | 1% |
| TsixStop_NPC | 975 | 806 | 806 | 0 | 83% | 83% | 0% |
| replicate F121_d0 | 393 | 49 | 27 | 22 | 12% | 7% | 6% |
| replicate F121_d7 | 47 | 17 | 17 | 0 | 36% | 36% | 0% |
| replicate F121_d20 | 89 | 66 | 44 | 22 | 74% | 49% | 25% |
| replicate TsixStop_d0 | 291 | 33 | 11 | 22 | 11% | 4% | 8% |
| replicate TsixStop_d7 | 51 | 16 | 11 | 5 | 31% | 22% | 10% |
| replicate TsixStop_d20 | 80 | 47 | 45 | 2 | 59% | 56% | 3% |

| dataset | XCI based on single cell expression | | | | | | |
|---|---|---|---|---|---|---|---|
| | total cells | XCI cells | 129 Xi cells | Cast Xi cells | XCI % | 129 Xi % | Cast Xi % |
| F121_d0 | 486 | 111 | 68 | 43 | 23% | 14% | 9% |
| F121_d3 | 127 | 57 | 31 | 26 | 45% | 24% | 20% |
| F121_d7 | 76 | 65 | 51 | 14 | 86% | 67% | 18% |
| F121_d11 | 143 | 116 | 85 | 31 | 81% | 59% | 22% |
| F121_NPC | 187 | 187 | 187 | 0 | 100% | 100% | 0% |

| dataset | XCI based on single cell accessibiity | | | | | | |
|---|---|---|---|---|---|---|---|
| | total cells | XCI cells | 129 Xi cells | Cast Xi cells | XCI % | 129 Xi % | Cast Xi % |
| F121_d0 | 164 | 42 | 31 | 11 | 26% | 19% | 7% |
| F121_d3 | 702 | 239 | 159 | 80 | 34% | 23% | 11% |
| F121_d7 | 77 | 61 | 42 | 19 | 79% | 55% | 25% |
| F121_d11 | 174 | 153 | 100 | 53 | 88% | 57% | 30% |
| F121_NPC | 178 | 170 | 169 | 1 | 96% | 95% | 1% |

The determination of XCI status by each modality is described in the Methods section. For the sci-Hi-C data, the long-range to mid-range difference (LMD) method was used on the allelic contact decay profiles (CDPs) of cells with at least 50 chr1 and chrX contacts. An LMD threshold value ($\delta_{Xi}^{LMD}$) of 0.9, 0.9, 0.88, and 0.95 was used for F121, ES_Tsix-stop, the F121 replicate experiment, and the ES_Tsix-stop replicate experiment, respectively. For the sci-RNA-seq data, the total allelic expression difference threshold ($\delta_{Xi}^{TAE}$) used was 1.21. For the sci-ATAC-seq data, the total allelic accessibility difference threshold ($\delta_{Xi}^{TAA}$) used was 0.98.

***Supplementary Table 3. sci-RNA-seq cell counts and expression summary statistics.***

| | XCI based on single cell interactions | | | | | | |
|---|---|---|---|---|---|---|---|
| **dataset** | **total cells** | **XCI cells** | **B6/129 Xi cells** | **Spret/Cast Xi cells** | **XCI %** | **B6/129 Xi %** | **Spret/Cast Xi %** |
| **F121_d0** | 986 | 116 | 59 | 57 | 12% | 6% | 6% |
| **F121_d3** | 435 | 61 | 34 | 27 | 14% | 8% | 6% |
| **F121_d7** | 243 | 107 | 68 | 39 | 44% | 28% | 16% |
| **F121_d11** | 164 | 82 | 52 | 30 | 50% | 32% | 18% |
| **F121_NPC** | 325 | 271 | 271 | 0 | 83% | 83% | 0% |
| **TsixStop_d0** | 89 | 11 | 8 | 3 | 12% | 9% | 3% |
| **TsixStop_d3** | 86 | 30 | 29 | 1 | 35% | 34% | 1% |
| **TsixStop_d7** | 159 | 103 | 100 | 3 | 65% | 63% | 2% |
| **TsixStop_d11** | 83 | 60 | 59 | 1 | 72% | 71% | 1% |
| **TsixStop_NPC** | 975 | 806 | 806 | 0 | 83% | 83% | 0% |
| **replicate F121_d0** | 393 | 49 | 27 | 22 | 12% | 7% | 6% |
| **replicate F121_d7** | 47 | 17 | 17 | 0 | 36% | 36% | 0% |
| **replicate F121_d20** | 89 | 66 | 44 | 22 | 74% | 49% | 25% |
| **replicate TsixStop_d0** | 291 | 33 | 11 | 22 | 11% | 4% | 8% |
| **replicate TsixStop_d7** | 51 | 16 | 11 | 5 | 31% | 22% | 10% |
| **replicate TsixStop_d20** | 80 | 47 | 45 | 2 | 59% | 56% | 3% |

| | XCI based on single cell expression | | | | | | |
|---|---|---|---|---|---|---|---|
| **dataset** | **total cells** | **XCI cells** | **129 Xi cells** | **Cast Xi cells** | **XCI %** | **129 Xi %** | **Cast Xi %** |
| **F121_d0** | 486 | 111 | 68 | 43 | 23% | 14% | 9% |
| **F121_d3** | 127 | 57 | 31 | 26 | 45% | 24% | 20% |
| **F121_d7** | 76 | 65 | 51 | 14 | 86% | 67% | 18% |
| **F121_d11** | 143 | 116 | 85 | 31 | 81% | 59% | 22% |
| **F121_NPC** | 187 | 187 | 187 | 0 | 100% | 100% | 0% |

| | XCI based on single cell accessibiity | | | | | | |
|---|---|---|---|---|---|---|---|
| **dataset** | **total cells** | **XCI cells** | **129 Xi cells** | **Cast Xi cells** | **XCI %** | **129 Xi %** | **Cast Xi %** |
| **F121_d0** | 164 | 42 | 31 | 11 | 26% | 19% | 7% |
| **F121_d3** | 702 | 239 | 159 | 80 | 34% | 23% | 11% |
| **F121_d7** | 77 | 61 | 42 | 19 | 79% | 55% | 25% |
| **F121_d11** | 174 | 153 | 100 | 53 | 88% | 57% | 30% |
| **F121_NPC** | 178 | 170 | 169 | 1 | 96% | 95% | 1% |

Counts of cells with at least 200 UMIs are given for each time point in the F121 sci-RNA-seq samples, along with mean and median UMIs per cell (see Extended Data Fig. 6A – C). Also provided are the number of cells with at least 160 UMIs per allele for F121 sci-RNA-seq samples, along with mean and median UMIs per allele per cell (see Extended Data Fig. 7A – D).

***Supplementary Table 4. sci-ATAC-seq cell counts and accessibility summary statistics.***

| | cells with >= 200 UMIs total expression | | | cells with >= 160 UMIs of expression per allele | | | | |
|---|---|---|---|---|---|---|---|---|
| **sample** | **cell count** | **mean UMIs** | **median UMIs** | **allelic cell counts** | **mean 129 UMIs** | **mean *cast* UMIs** | **median 129 UMIs** | **median *cast* UMIs** |
| **F121_d0** | 1,682 | 1,941 | 1,562 | 540 | 302 | 291 | 265 | 258 |
| **F121_d3** | 868 | 1,179 | 910 | 142 | 288 | 281 | 244 | 245 |
| **F121_d7** | 501 | 1,324 | 947 | 104 | 283 | 279 | 254 | 248 |
| **F121_d11** | 574 | 1,793 | 1,421 | 214 | 312 | 309 | 272 | 277 |
| **F121_NPC** | 497 | 2,509 | 2,158 | 344 | 344 | 342 | 302 | 318 |
| **F123_d0** | 1,654 | 1,923 | 1,538 | 503 | 295 | 275 | 263 | 240 |
| **F123_d3** | 646 | 902 | 626 | 73 | 295 | 265 | 273 | 247 |
| **F123_d7** | 1,102 | 1,210 | 915 | 181 | 276 | 253 | 250 | 221 |
| **F123_d11** | 925 | 1,415 | 1,028 | 206 | 302 | 279 | 264 | 242 |
| **F123_NPC** | 572 | 2,536 | 2,254 | 389 | 366 | 330 | 335 | 302 |

Counts of cells with at least 500 UMIs are given for each time point in the F121 sci-ATAC-seq samples, along with mean and median UMIs per cell (see Extended Data Fig. 6D – F). Also provided are the number of cells with at least 500 UMIs per allele for F121 sci-ATAC-seq samples, along with mean and median UMIs per allele per cell (see Extended Data Fig. 7E – H).

***Supplementary Table 5. MMD-MA parameters, cell downsampling and results.***

| | | MMD-MA parameters | | | AUC scores | | | | | |
|---|---|---|---|---|---|---|---|---|---|---|
| | | features | lambda 1 | lambda 2 | d0 | d3 | d7 | d11 | NPCs | mean |
| **sci-RNA-seq vs sci-ATAC-seq** | genome-wide | 6 | 10^-3 | 10^-5 | 0.972 | 0.832 | 0.829 | 0.850 | 1.000 | 0.907 |
| | autosomes | 4 | 10^-3 | 10^-4 | 0.974 | 0.840 | 0.821 | 0.865 | 1.000 | 0.911 |
| | chrX | 3 | 10^-3 | 10^-5 | 0.677 | 0.547 | 0.516 | 0.499 | 0.905 | 0.650 |
| | chr1 | 3 | 10^-7 | 10^-3 | 0.872 | 0.773 | 0.723 | 0.779 | 0.998 | 0.848 |
| | chr1+chrX | 3 | 10^-3 | 10^-5 | 0.904 | 0.750 | 0.732 | 0.821 | 0.999 | 0.863 |
| **sci-RNA-seq vs sci-Hi-C** | genome-wide | 5 | 10^-6 | 10^-7 | 0.779 | 0.694 | 0.717 | 0.727 | 0.788 | 0.759 |
| | autosomes | 3 | 10^-4 | 10^-7 | 0.687 | 0.727 | 0.738 | 0.673 | 0.616 | 0.684 |
| | chrX | 3 | 10^-4 | 10^-3 | 0.802 | 0.601 | 0.569 | 0.467 | 0.820 | 0.728 |
| | chr1 | 3 | 10^-3 | 10^-3 | 0.761 | 0.526 | 0.441 | 0.314 | 0.699 | 0.648 |
| | chr1+chrX | 3 | 10^-7 | 10^-5 | 0.773 | 0.526 | 0.647 | 0.654 | 0.834 | 0.732 |
| **sci-ATAC-seq vs sci-Hi-C** | genome-wide | 3 | 10^-6 | 10^-6 | 0.773 | 0.699 | 0.621 | 0.617 | 0.639 | 0.683 |
| | chrX | 5 | 10^-4 | 10^-3 | 0.661 | 0.629 | 0.540 | 0.527 | 0.836 | 0.662 |

| **sci-RNA-seq vs sci-ATAC-seq** | sci-RNA-seq | | | | | | sci-ATAC-seq | | | | | |
|---|---|---|---|---|---|---|---|---|---|---|---|---|
| | d0 | d3 | d7 | d11 | NPC | total | d0 | d3 | d7 | d11 | NPC | total |
| original | 404 | 98 | 73 | 164 | 285 | 1,024 | 165 | 705 | 77 | 177 | 180 | 1,304 |
| subsampled | 129 | 98 | 60 | 138 | 141 | 566 | 165 | 124 | 77 | 177 | 180 | 723 |
| subsampled proportion | 23% | 17% | 11% | 24% | 25% | | 23% | 17% | 11% | 24% | 25% | |

| **sci-RNA-seq vs sci-Hi-C** | sci-RNA-seq | | | | | | sci-Hi-C | | | | | |
|---|---|---|---|---|---|---|---|---|---|---|---|---|
| | d0 | d3 | d7 | d11 | NPC | total | d0 | d3 | d7 | d11 | NPC | total |
| original | 404 | 98 | 73 | 164 | 285 | 1,024 | 986 | 435 | 243 | 164 | 325 | 2,153 |
| subsampled | 404 | 98 | 73 | 78 | 154 | 807 | 849 | 206 | 153 | 164 | 325 | 1,697 |
| subsampled proportion | 50% | 12% | 9% | 10% | 19% | | 50% | 12% | 9% | 10% | 19% | |

| **sci-ATAC-seq vs sci-Hi-C** | sci-ATAC-seq | | | | | | sci-Hi-C | | | | | |
|---|---|---|---|---|---|---|---|---|---|---|---|---|
| | d0 | d3 | d7 | d11 | NPC | total | d0 | d3 | d7 | d11 | NPC | total |
| original | 165 | 705 | 77 | 177 | 180 | 1,304 | 986 | 435 | 243 | 164 | 325 | 2,153 |
| subsampled | 165 | 263 | 77 | 99 | 180 | 784 | 272 | 435 | 127 | 164 | 297 | 1,295 |
| subsampled proportion | 21% | 34% | 10% | 13% | 23% | | 21% | 34% | 10% | 13% | 23% | |

MMD-MA parameters and area under the curve (AUC) scores for all the pairwise alignments reported in the study, along with the number of cells before and and after subsampling the cells from the five time-points (d0, d3, d7, d11, NPCs) to ensure that the fraction of cells in each time-point was consistent across the two datasets (see Methods).

### Extended Data Figures

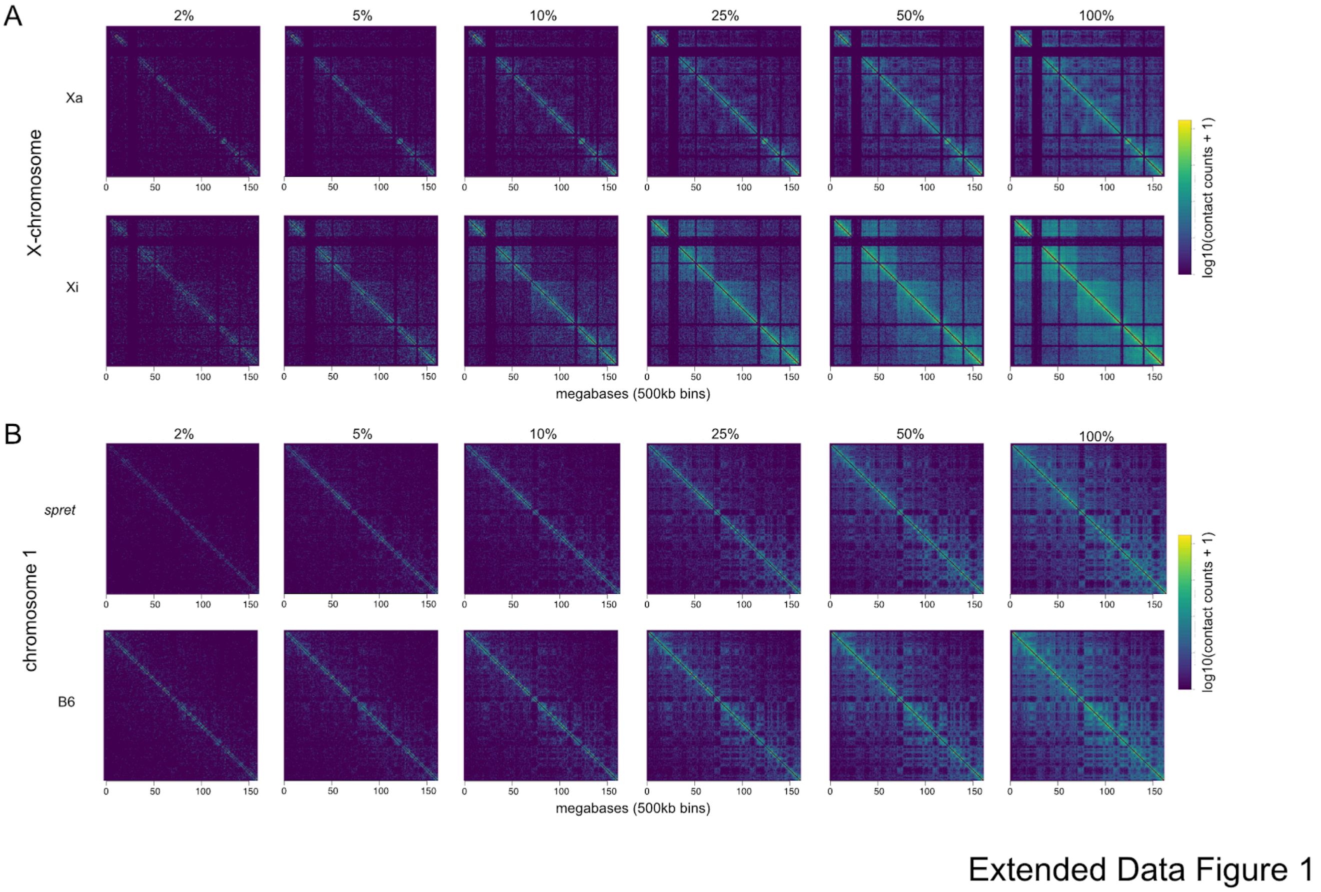


***Extended Data Fig. 1. Exemplar pseudobulk sci-Hi-C contact maps using different numbers of Patski cells.***

**A)** Aggregate (pseudobulk) allelic contact maps of Xi and Xa using 2-100% (4,493 total cells) of randomly sampled available cells from the Patski sci-Hi-C experiment. Color scale reflects log contact counts.

**B)** As in A, but for maternal (B6) and paternal (*spret)* alleles of chr1.

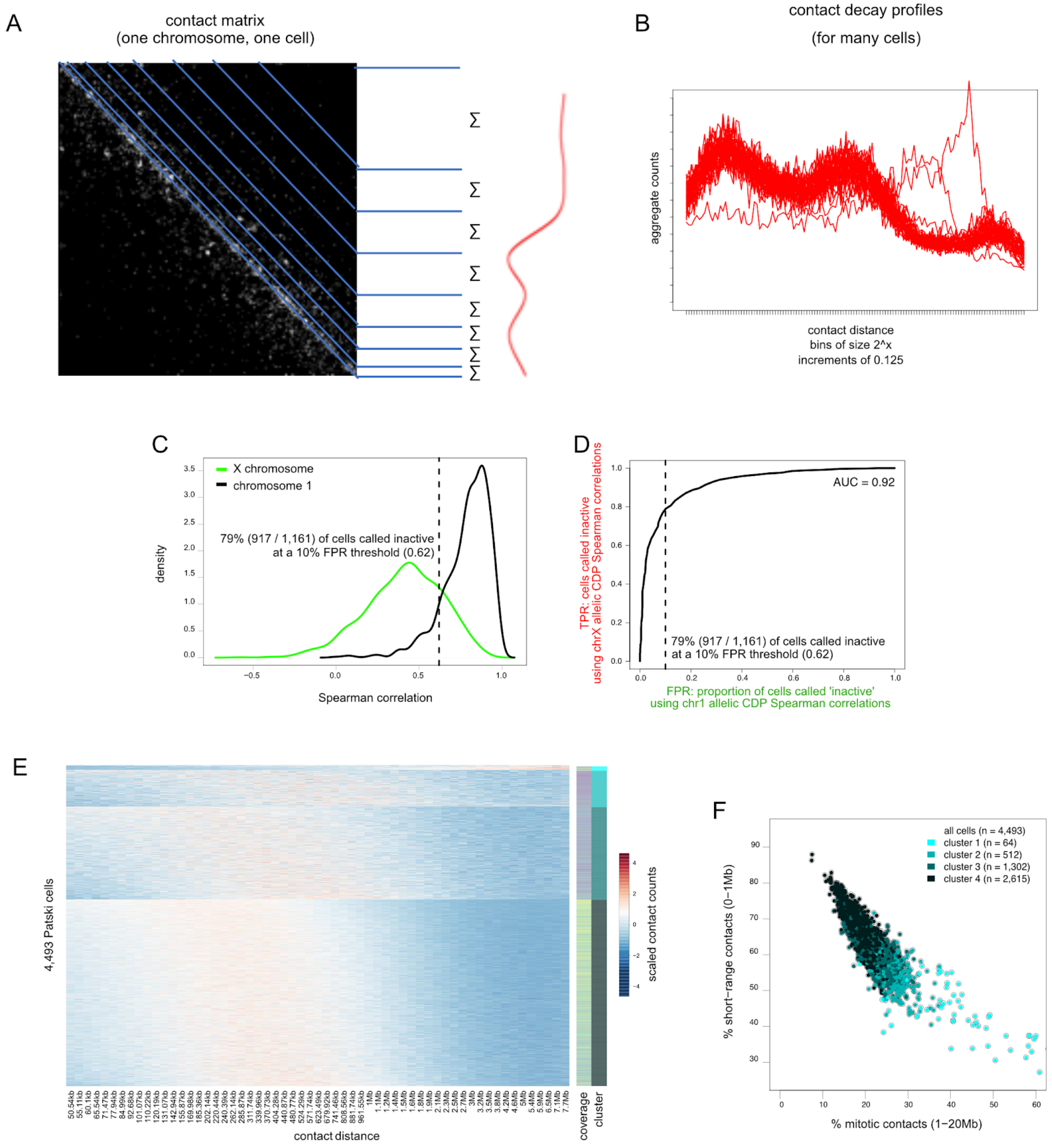


Extended Data Figure 2

***Extended Data Fig. 2. Strategy to determine CDPs and cell cycle state by sci-Hi-C.***

**A)** Example of a contact map for a single chromosome of a single cell based on sci-Hi-C data overlaid with a schematic of exponentially-increasing contact distances ranges used to summate contact counts with respect to distance in order to produce contact decay profiles (CDPs), shown to the right.

**B)** Examples of CDPs for a single chromosome in many single cells.

**C)** Distributions of Spearman correlations between the CDPs of each allele for each of 1,161 cells for chrX (green; as described in Fig. 2B) and chr1 (black; as described in Fig. 2F). The dashed line represents a threshold chosen below which cells were called as having a bipartite Xi. This threshold represents a 10% false positive rate (FPR) based on the distribution allelic CDP Spearman correlations for chr1 (see D).

**D)** Plot of true positive rate (TPR; proportion of Patski cells called as having a bipartite Xi based on the Spearman correlations between allelic CDPs for chrX as described in Fig. 2B, C) versus false positive rate (FPR; proportion of Patski cells called 'inactive' based on the Spearman correlations between allelic CDPs for chr1 as described in Fig. 2F, C) at all thresholds of Spearman correlation value (0 to 1). The area under the curve (AUC) is 0.92. The dashed line represents the 10% FPR threshold chosen below which cells were called as having a bipartite Xi based on chrX allelic CDP Spearman correlation.
**E)** Heatmaps of biallelic CDPs for the autosomes of 4,493 Patski cells clustered into four groups using *k*-means clustering and the Spearman correlation distance between the cells' CDPs. Heatmap colors reflect z-scaled counts within logarithmic bins with the left-hand boundaries specified along the bottom from 50 kb to 8 Mb.
**F)** Scatter plot of the proportion of short-range contact pairs (within 1 Mb) versus mitotic contact pairs (between 1 and 20 Mb) for 4,493 Patski cells. Cells colored by the *k*-means clusters described in C**.**

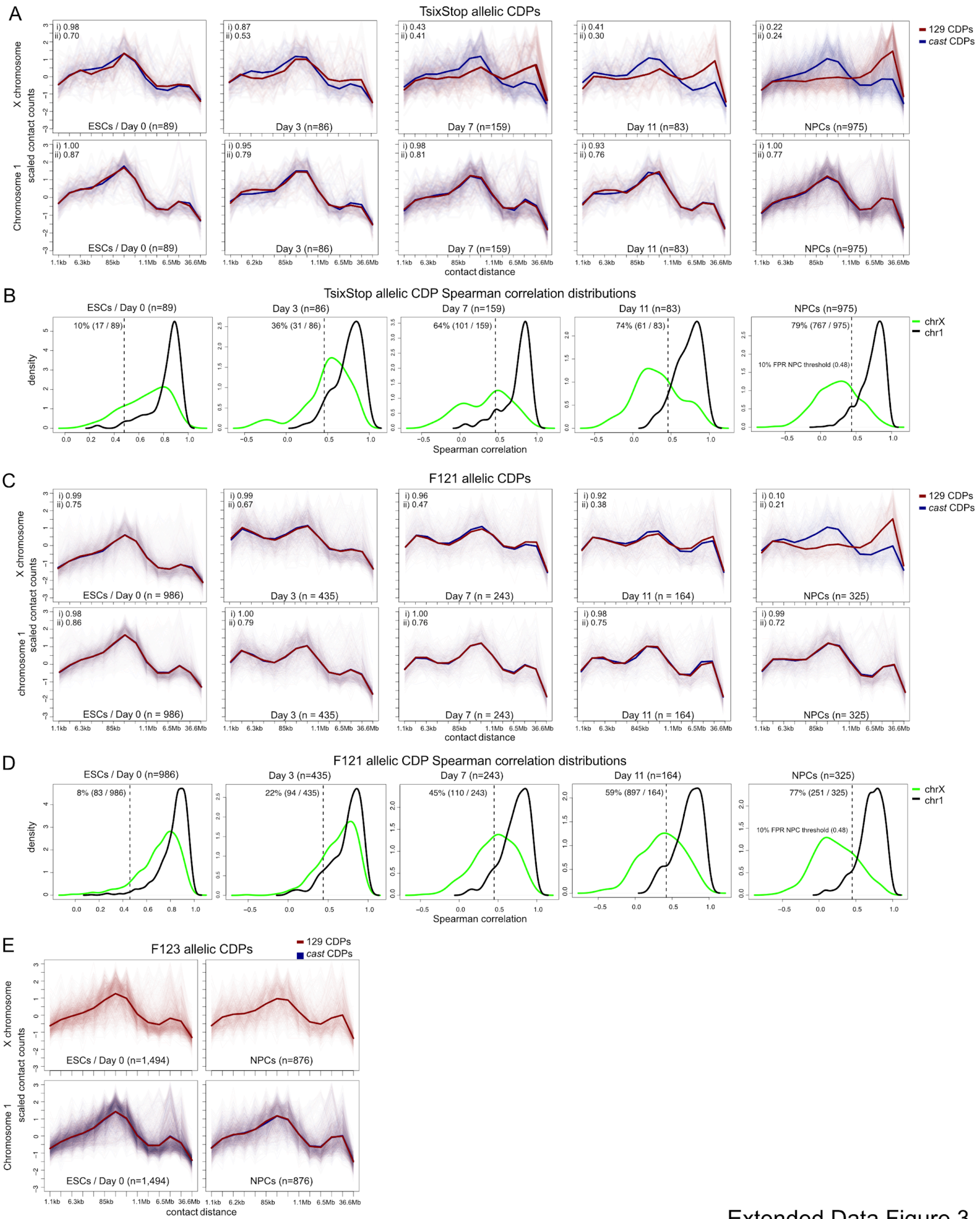


Extended Data Figure 3

***Extended Data Fig. 3. CDPs in ES_Tsix-stop, F121, and F123 cells during differentiation.***
**A)** Plots of allelic contact decay profiles (CDPs) for ES_Tsix-stop cells with at least 50 contacts per allele within chrX (top row; Xi, red; Xa, blue) and chr1 (bottom row; *cast*, red; 129, blue) for the indicated time points during differentiation from ESCs (d0) to EBs (day 11), as well as differentiated NPCs. The plots show z-scaled counts within contact distance ranges shown along the x-axis. Contact counts were binned within exponentially-increasing contact distances ranges (2^x where x was incremented by 0.125). These bins were then aggregated further to reduce noise by combining the counts within 10 non-overlapping bins at a time. The number of cells at each time point is given by the n number. The average allelic CDP is plotted over the plots of CDPs for each individual cell. Spearman correlation values between the two mean allelic CDPs (i) as well as the median of the Spearman correlation values between the allelic CDPs for each individual cell (ii) are given in the top left corner of each plot.
**B)** Distributions of Spearman correlations between CDPs of each allele for each of the cells as in A for chrX (green) and chr1 (black). The dashed line represents a threshold chosen below which cells were called having a bipartite Xi based on the chrX allelic CDP Spearman correlation. This threshold represents a 10% false positive rate (FPR) based on the distribution for chr1 allelic CDP Spearman correlations.
**C)** As in A, but for F121 cells.
**D)** As in B, but for F121 cells.
**E)** As in A, but for F123 ESCs and NPCs. Note that these male cells only possess a 129 chrX.

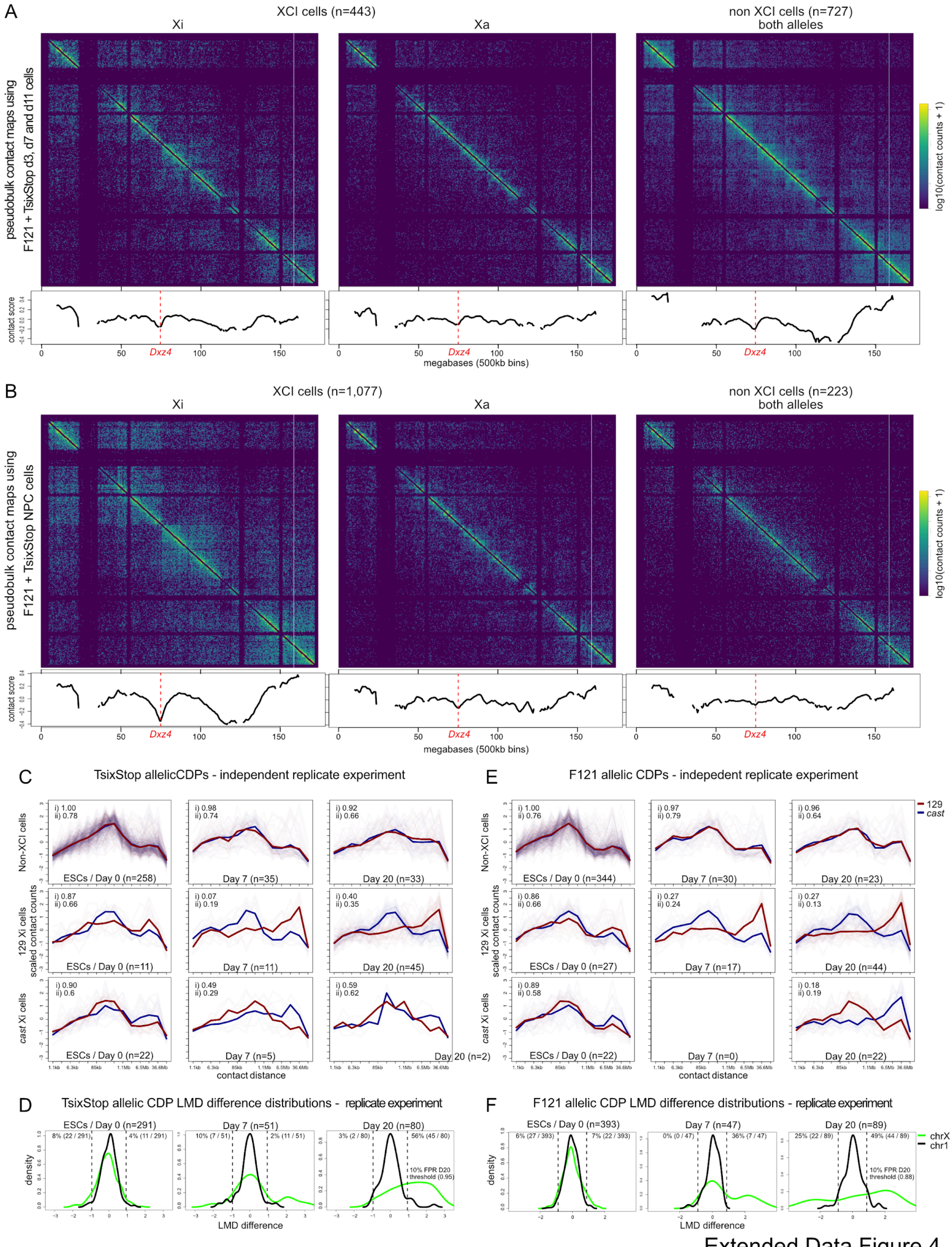


Extended Data Figure 4

***Extended Data Fig. 4. EB and NPC allelic contact maps and replicate CDPs in ES_Tsix-stop and F121.***
**A)** Aggregate allelic contact maps combining the cells that were classified as having a chrX homolog exhibiting an Xi bipartite structure (XCI cells) and those that did not (non-XCI cells) for EB cells (d3, d7, and d11 cells from both F121 and ES_Tsix-stop time course experiments combined). Alleles were combined for non-XCI cells. Below each contact map is a plot of the contact score (see Methods) showing the dip in contacts across the *Dxz4* locus (vertical red dashed line).
**B)** As in A, but for NPCs from both the F121 and ES_Tsix-stop time course experiments.
**C)** Plots of allelic CDPs for ES_Tsix-stop for the indicated time points during differentiation from ESCs to d20 (beyond embryoid bodies) for cells classified as not having a bipartite Xi (top row), and cells that do exhibit CDPs indicative of an bipartite Xi on their 129 allele (middle row; XCI is skewed towards the 129 allele) or their *cast* allele (bottom row; XCI is skewed towards the *cast* allele). Only cells with at least 50 contacts per allele within both chrX and chr1. The plots show z-scaled counts within contact distance ranges shown along the x-axis. Contact counts were binned within exponentially-increasing contact distances ranges (2^x where x was incremented by 0.125). These bins were then aggregated further to reduce noise by combining the counts within 10 non-overlapping bins at a time. The number of cells at each time point is given by the n number. The average allelic CDP is plotted over the plots of CDPs for each individual cell. Spearman correlation values between the two mean allelic CDPs (i) as well as the median of the Spearman correlation values between the allelic CDPs for each individual cell (ii) are given in the top left corner of each plot.
**D)** Distributions of the difference between the long-range to mid-range differences (LMDs) of each allele (Fig. 2I and Methods) for each of the cells as described in C for chrX (green) and chr1 (black). The dashed lines represent a threshold chosen below which cells were called as having a bipartite Xi based on chrX allelic CDP LMD differences. This threshold represents a 10% false positive rate (FPR) based on the distribution for chr1 allelic CDP LMD differences.
**E)** As in C, but for F121 cells.
**F)** As in D, but for F121 cells.

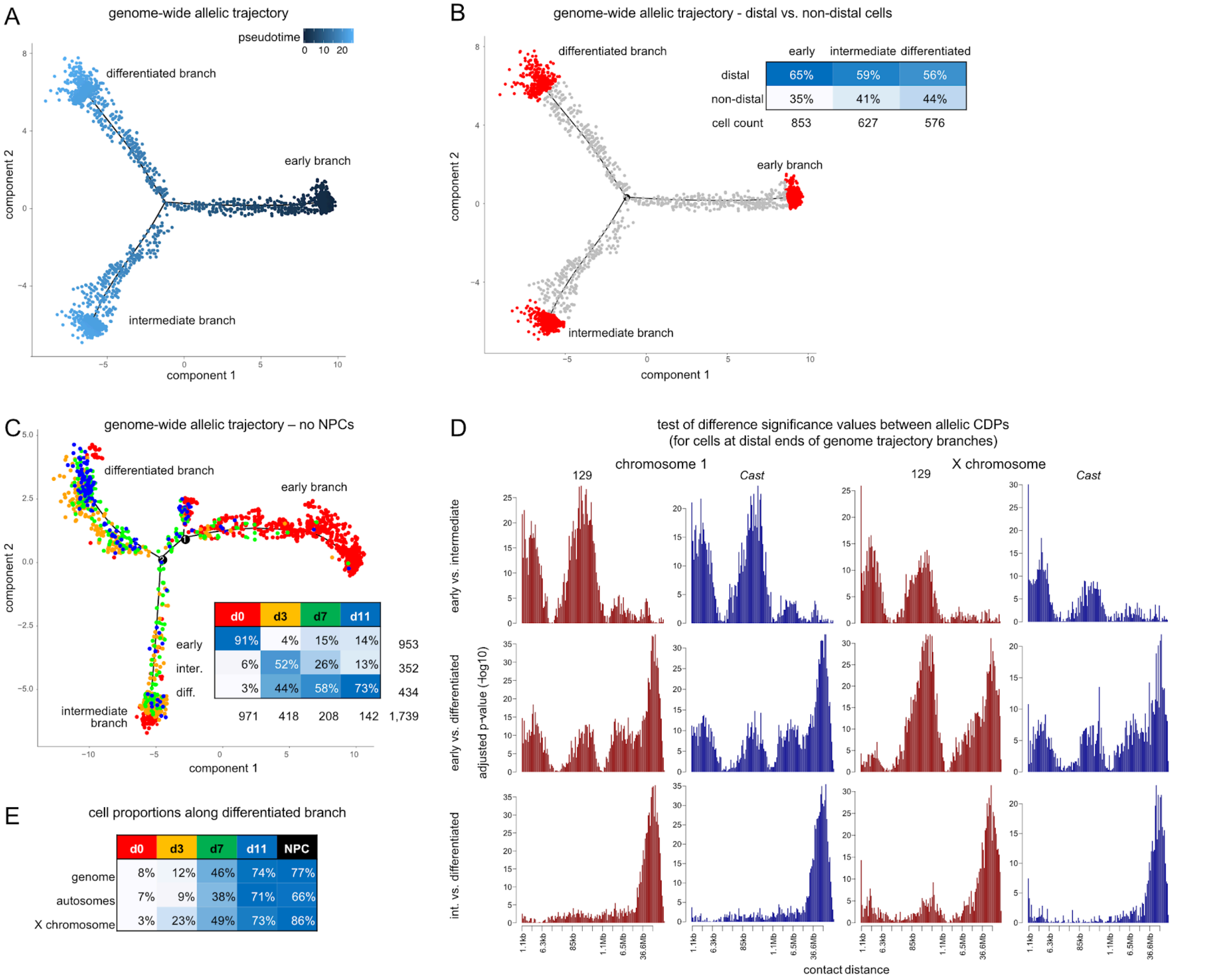


Extended Data Figure 5

***Extended Data Fig. 5. Trajectory analysis of F121 cells based on allelic CDPs.***

**A)** As in Fig. 4A, but cells colored by pseudotime with respect to the root ESC branch.

**B)** As in Fig. 4A, but highlighting the most distal cells on each branch of the trajectory. The inset table gives the proportion of cells classified as distal and non-distal. Pseudotimes for each branch were renormalized to fall between 0 and 1 and the following thresholds were used to classify cells as distal for the early, intermediate, and differentiated branches respectively: 0.91, 0.83, and 0.73.

**C)** As in Fig. 4A, but excluding NPCs.

**D)** Plots of the -log10 adjusted p-values from t-tests between the values within every logarithmic bin (2^x where x was incremented by 0.125) of the allelic chr1 and chrX CDPs (129, red; *cast,* blue) for cells at the distal ends of each pair of branches (rows) in the trajectory described in Fig. 4A (see B).

**E)** Table of the percentage of cells per time point along the differentiated branch of the genome-wide, autosomal and X-chromosome trajectories (as in Fig. 4A – C).

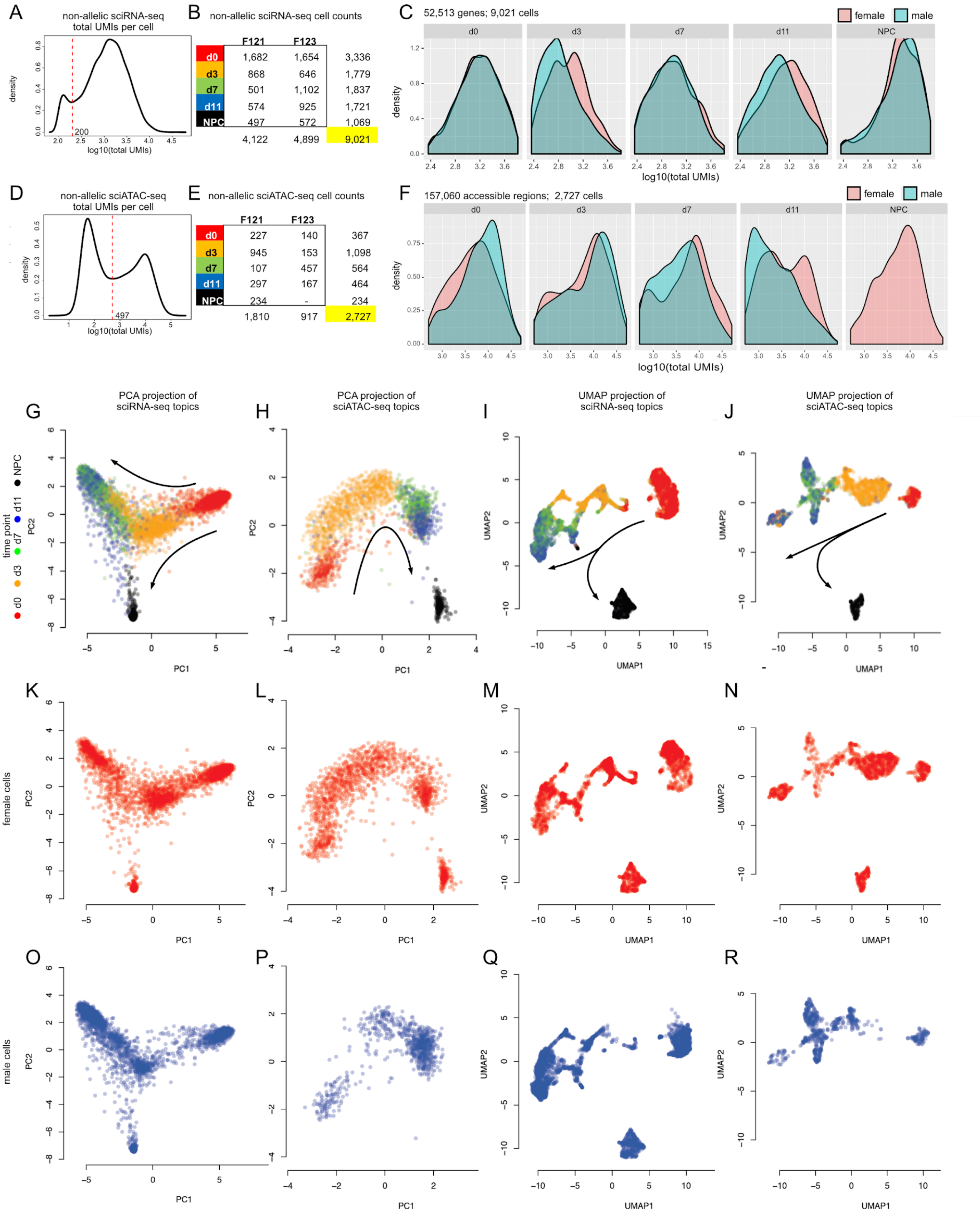


Extended Data Figure 6

***Extended Data Fig. 6. Sex differences in non-allelic expression and accessibility data distribution, and topic modeling projections.***

**A)** Distributions of total genome-wide expression (log10[UMIs]) from sci-RNA-seq in F121 and F123 cells. The vertical line shows the ~200 UMI minima used as a threshold to filter out cells with low expression.
**B)** Table of F121 and F123 cell counts per time point after applying the filter described in A.
**C)** Distributions of total genome-wide expression (log10[UMIs]) from sci-RNA-seq in F121 and F123 cells faceted by time point in cells showing a minimum total expression of 200 UMI.
**D)** As in A, but based on total genome-wide chromatin accessibility (log10[UMIs]) from sci-ATAC-seq in F121 and F123 cells. The vertical line shows the ~500 UMI minima used as a threshold to filter out cells with low values.
**E)** As in B, but based on total genome-wide accessibility (log10[UMIs]) from sci-ATAC-seq in F121 and F123 cells.
**F)** Distributions of total genome-wide accessibility (log10[UMIs]) from sci-ATAC-seq in F121 and F123 cells faceted by time point in cells showing a minimum total accessibility counts of 500 UMI.
**G)** Projection using the first two components from PCA of a topic matrix generated from non-allelic F121 and F123 sci-RNA-seq data. The matrix of topics per cell was decomposed from a matrix of UMI counts within genes for each cell. Each data point represents a cell colored by time point. Arrows indicate the direction of development. (The topic matrix served as input to the MMD-MA algorithm - see section 5).
**H)** As in G, but for non-allelic sci-ATAC-seq data. The matrix of topics per cell was decomposed from a matrix of UMI counts within accessible regions for each cell.
**I)** As in G, but using a UMAP projection.
**J)** As in H, but using a UMAP projection.
**K-N)** As in G-J, but for F121 cells only.
**O-R)** As in G-J, but for F123 cells only.

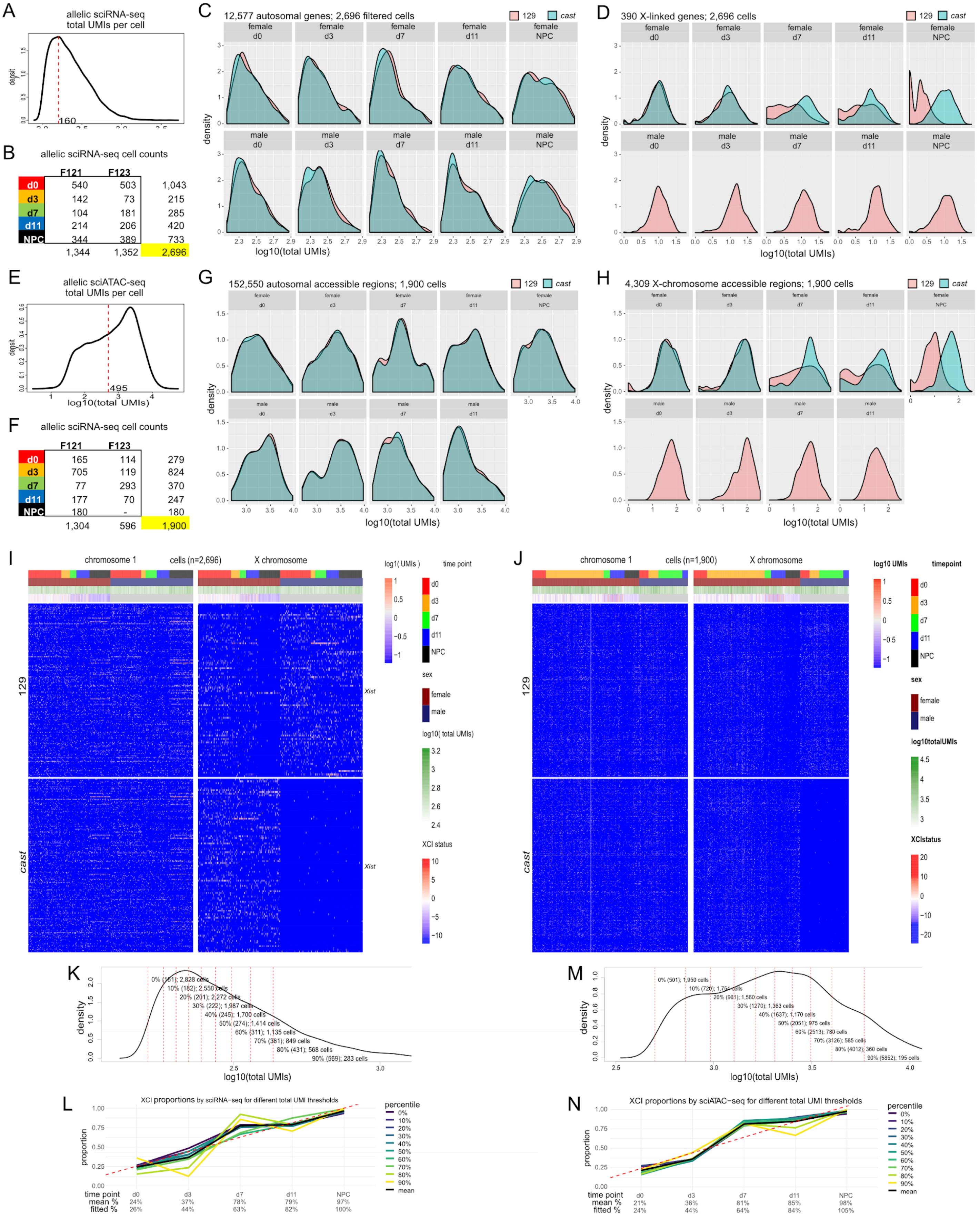


Extended Data Figure 7

***Extended Data Fig. 7. Allelic expression and accessibility distributions, filtering, and visualization.***
**A)** Distributions of total genome-wide allelic expression (log10[UMIs]) from sci-RNA-seq in F121 and F123 cells. The vertical line shows the ~160 UMI maxima used as a threshold to filter out cells with low expression. This value was used since no minima was available in the distribution to use as a threshold.
**B)** Table of F121 and F123 cell counts per time point after applying the filter described in A.
**C)** Allelic expression (log10[UMIs]) distributions based on sci-RNA-seq data for autosomal genes faceted by time point (columns) and sex (rows).
**D)** As in C, but for expression distributions based on X-linked genes.
**E)** As in A, but based on total genome-wide accessibility (log10[UMIs]) from sci-ATAC-seq in F121 and F123 cells. The vertical line shows the ~500 UMI value at an inflection point in the distribution used as a threshold to filter out cells with low expression. This value was used since no minima was available in the distribution to use as a threshold and it closely matched the value used for non-allelic data.
**F)** As in B, but based on total genome-wide allelic accessibility (log10[UMIs]) from sci-ATAC-seq on F121 and F123 cells.
**G)** As in C, but for accessibility based on allelic sci-ATAC-seq data.
**H)** As in G, but for accessibility distributions based on accessible regions along chrX.
**I)** Heatmap of allelic expression (log UMIs) as measured by sci-RNA-seq for chr1 (left) and chrX (right) for the 25% most variability expressed genes per chromosome (rows) for 2,696 F121 and F123 cells that passed filter. Cells sorted by annotations described in legend on the right of the heat maps.
**J)** As in I, but for allelic accessibility as measured by sci-ATAC-seq 25% most variability accessible regions per chromosome (rows) for 1,900 F121 and F123 cells that passed filter.
**K)** Distributions of total allelic UMIs per cell for sci-RNA-seq. Vertical lines represent quantile thresholds used in robustness testing in B.
**L)** Plots of the proportion (%) of F121 cells with a silenced chrX using sci-RNA-seq expression and the method described in Fig. 6C for different proportions of cells based on total expression quantiles. This shows that the proportions of cells showing XCI silencing based on allelic sci-RNA-seq is robust to coverage level and cell numbers.
**M)** As in K, but based on sci-ATAC-seq accessibility data.
**N)** Plots of the proportion (%) of F121 cells with a low accessibility X chromosome data using sci-ATAC-seq accessibility data and the method described in Fig. 6D for different proportions of cells based on total accessibility quantiles. This shows that the proportions of cells that have undergone XCI based on allelic sci-ATAC-seq is robust to coverage level and cell numbers.

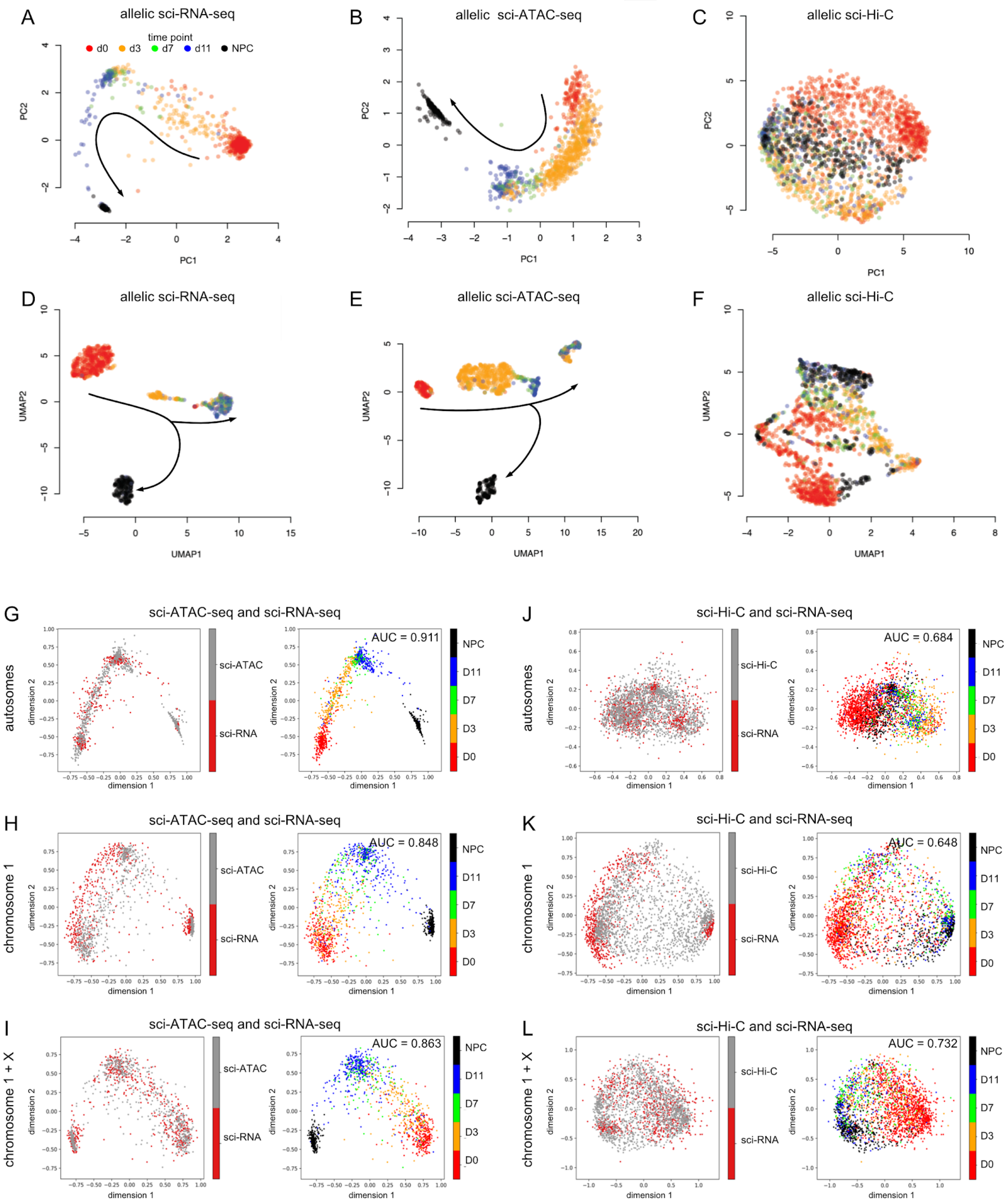


Extended Data Figure 8

***Extended Data Fig. 8. Projections of allelic topic matrices used for multimodal alignment.***
**A)** Projection using the first two components from PCA of a topic matrix generated from allelic sci-RNA-seq data that served as input to the MMD-MA algorithm. The matrix of topics per cell was decomposed from a matrix of UMI counts within genes from each allele concatenated together (rows) for each cell (columns). Each data point represents a cell colored by time point. Arrows indicate the direction of development.
**B)** As in A, but for allelic sci-ATAC-seq data. The matrix of topics per cell was decomposed from a matrix of UMI counts within accessible regions from each allele concatenated together (rows) for each cell (columns).
**C)** As in A, but for allelic sci-Hi-C data. The matrix of topics per cell was decomposed from a matrix of concatenated contact decay counts within logarithmically-increasing sized bins for both alleles of each chromosome (rows) for each cell (columns).
**D)** As in A, but using a UMAP projection.
**E)** As in B, but using a UMAP projection.
**F)** As in C, but using a UMAP projection.
**G)** As in Figure 7A, but only using autosomal data.
**H)** As in Figure 7A, but only using chr1 data.
**I)** As in Figure 7A, but only using data from chr1 and chrX.
**J)** As in Figure 7B, but only using autosomal data.
**K)** As in Figure 7B, but only using chr1 data.
**L)** As in Figure 7B, but only using data from chr1 and chrX.

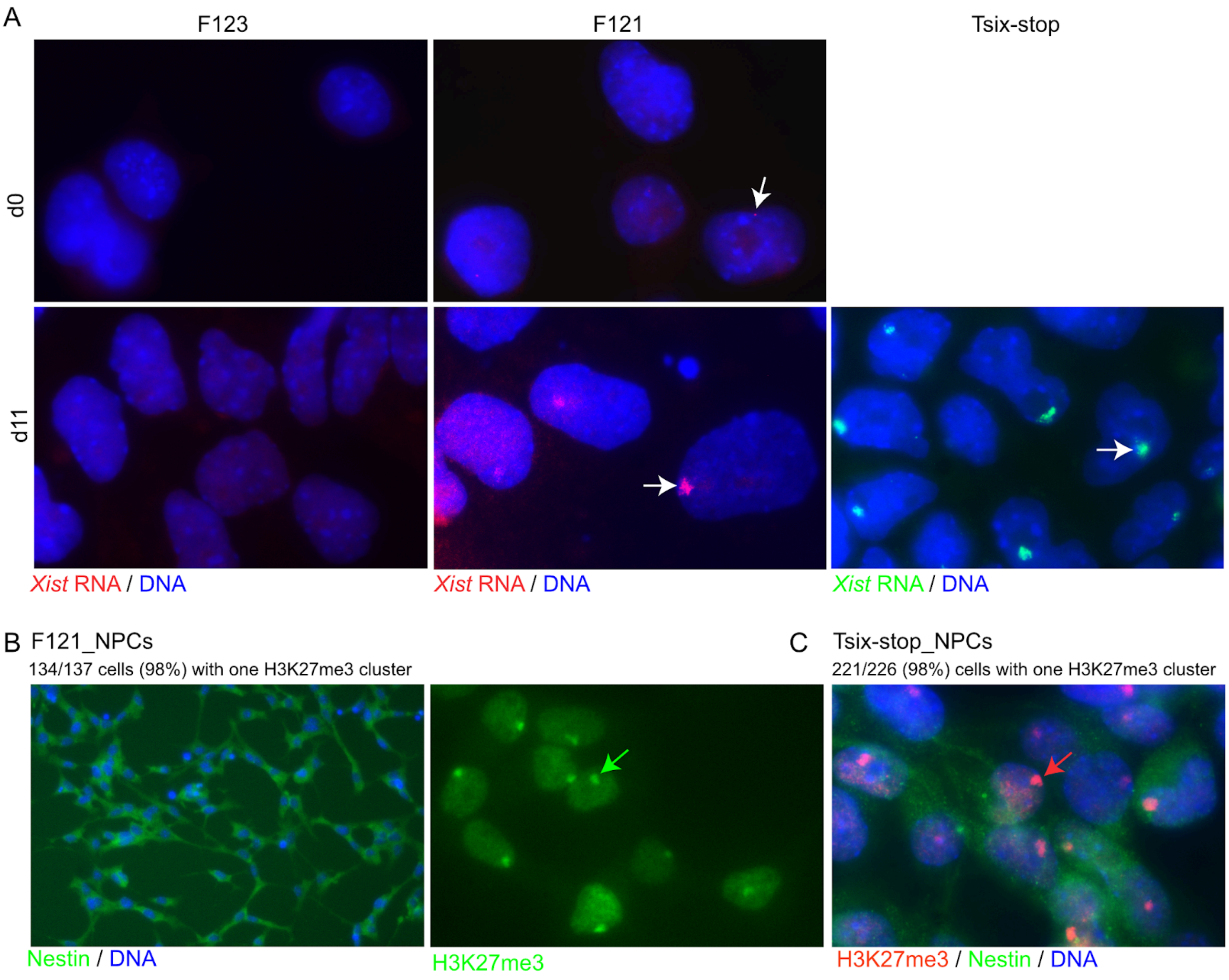


Extended Data Figure 9

***Extended Data Fig. 9. Verification of establishment of XCI during in vitro differentiation of mESCs.***
**A)** RNA-FISH using an *Xist* probe in male and female d0 and d11 cells show the presence of an *Xist* cloud (arrows) in d11 differentiated cells from the female lines F121 and ES_Tsix-stop, but not the male F123 cells or d0 F121 female ESCs. 91/96 (95%) of ES_Tsix-stop d11 cells show one *Xist* cloud, indicating the establishment of XCI. Note that some d0 cells show pinpoints of signals (arrowhead), indicating low levels of *Xist* expression in undifferentiated mESCs. *Xist* RNA-FISH was done using green or red fluorescence-labeled probes using a 10 kb *Xist* cDNA plasmid (pXho, which contains most of *Xist* exon 1). DNA is stained by Hoechst 33258 (blue).
**B)** Immunostaining using a H3K27me3 antibody detected by a FITC-conjugated secondary antibody (green) was done to mark the Xi in NPCs differentiated from F121. DNA is stained by Hoechst 33258 (blue). Immunostaining for nestin with an Alexa Fluor 488-conjugated antibody (green) confirmed differentiation into NPCs. 134/137 (98%) NPCs were scored as having one H3K27me3 cluster, indicating the establishment of XCI.
**C)** Co-immunostaining with a H3K27me3 antibody detected by a Texas red-conjugated secondary antibody (red) to mark the Xi and an Alexa Fluor 488-conjugated antibody for nestin (green) was done in ES_Tsix-stop NPCs. DNA is stained by Hoechst 33258 (blue). 221/226 (98%) NPCs were scored as having one H3K27me3 cluster, indicating the establishment of XCI.

### Extended Data Video

***Extended Data Video 1. Patski cell Xi bipartite structure re-emerges after mitosis.***
A movie compiled from aggregate allelic contact maps for the Xi and Xa (top row) and B6 and *spretus* chromosome 1 homologs (bottom row) using contact pairs from 64 mitotic, 512 early interphase, 1,032 mid interphase, and 2,615 late interphase cells, as grouped using the k-means clustering of the autosomal CDPs (Extended Data Fig. 3E).
